# MAFA drives pancreatic islet maturation through coordination of cell growth, metabolic maturation, and resilience to stress

**DOI:** 10.64898/2026.09.29.755533

**Authors:** Veronica A. Cochrane, Michael Sterr, Marianne Aniag, Sara Jimenez, Madeline S. Wang, Ying Liu, Alena Boos, Juliane Merl-Pham, Aarti Anand, Ryan E. Hoang, Rachita Gupta, Heiko Lickert, Tiago C. Alves, Jeeyeon Cha, Roland Stein, Fabian J. Theis, Rafael Arrojo e Drigo, Audrey V. Parent, Matthias Hebrok

## Abstract

Stem cell-derived islets (SC-islets) are a promising source for replacing the loss of insulin-secreting β cells in insulin-dependent diabetes patients, but compared to their primary human counter parts, SC-islets are considered functionally immature and do not immediately regulate blood glucose after transplantation. To better understand the molecular underpinnings of β cell maturation, we genetically engineered human pluripotent stem cells (hPSCs) to control the induction of the β cell-maturing transcription factor MAFA or a MAFA^S64F^ mutant variant associated with premature β cell aging and senescence in SC-islets. We employ a combination of *in vitro* assays, multi-omics analyses, and murine *in vivo* transplantation studies to demonstrate that MAFA remodeling of the SC-islet transcriptional landscape promotes SC-islet biogenesis and metabolic maturation giving rise to greater SC-islet resilience and insulin secretory capacity. Parallel MAFA^S64F^ studies highlight the importance for compensatory mechanisms for sustaining accelerated oxidative metabolism and enhanced insulin secretion without inducing intracellular stress responses and islet dysfunction. Overall, these results reveal MAFA-governed transcriptional changes that drive SC-islet functional maturation, which provides the foundation for maturing SC-islets *in vitro*, preserving β cell integrity, and improving SC-islet cell replacement therapies.

## Introduction

The directed differentiation of hPSCs to stem cell-derived islets (SC-islets) containing insulin-producing SC-β cells has emerged as a promising cell replacement therapy for treating insulin-dependent diabetes. Notably, *in vitro* assessment of SC-islets pre-transplantation repeatedly demonstrates that they do not fully recapitulate endogenous human islets with regards to their transcriptome, metabolism, and bi-phasic secretory capacity^1–7^. However, after transplantation in murine models and Type 1 Diabetes patient recipients, SC-islets undergo maturation over ∼6 months wherein SC-islets attain greater insulin content and regulated secretory capacity improving their ability to maintain blood glucose homeostasis^1,4–6,8^. Thus, increasing our understanding of the processes that contribute to β cell maturation is essential for timely β cell functionality post-transplantation and improving the outcome of cell replacement therapies.

A key β cell transcription factor that has been implicated in establishing adult β cell functionality is MAFA. MAFA expression has been found to be increased during human adolescent years (≥9yo)^9,10^. Adolescence is a period of rapid physical growth, and the increased metabolic demands of the body are accompanied by insulin resistance. This marks a pivotal time in β cell development wherein β cells increase insulin production and secretion to compensate for the greater insulin demand^11^. In addition to adolescent β cell development, MAFA expression has also been found to correlate with SC-islet functionality wherein SC-islets differentiated *in vitro* have low functionality and low MAFA expression, but there is a correlative increase of both several months post-transplantation *in vivo*^3–5,12^. The role of MAFA in β cell maturation is of particular interest because MAFA has been shown to target insulin expression, β cell identity markers and insulin secretory mechanisms^9,13–16^.

Perturbations of MAFA activity has been associated with β cell dysfunction in both type 2 diabetes and maturity onset diabetes of the youth (MODY). MAFA expression has been found to be reduced in β cells from human type 2 diabetes patients and mouse models of type 2 diabetes^17^. Loss of MAFA has been tied to reduced expression of insulin, β cell identity markers, and genes involved in glucose sensing and insulin secretion, which can be ameliorated by overexpressing MAFA in β cells of type 2 diabetes mouse models^18^. Conversely, a MAFA^S64F^ mutant variant exhibiting stabilization and hyperactivity has been identified as a MODY gene^19^. MafA^S64F^ mouse models recapitulating the human phenotype have found that the β cells undergo a period of hyperactivity before becoming non-functional and prematurely senescent^20^. These findings indicate that MAFA activity must be tightly regulated for optimal β cell function.

Although MAFA has been extensively studied in mice, we still lack a comprehensive understanding of the cellular processes that it governs in human model systems. In the present study, we induce MAFA expression in human SC-islets to mimic adolescent β cell maturation and identify key biological processes and cellular pathways that give rise to enhanced insulin secretory capacity and islet resilience.

## Results

### MAFA^WT^ overexpression increases SC-islet β-cell composition and insulin secretory capacity

Stem cell differentiation into SC-islets offer a unique opportunity to investigate MAFA’s function in a human relevant β cell model. SC-β cells indeed more closely resemble immature β cells in terms of their insulin secretion, mitochondrial metabolic activity, and transcriptome^1–6^. This has in part been attributed to SC-β cells expressing low or undetectable levels of MAFA^1–5,7^. We thus implemented a doxycycline (dox)-inducible MAFA overexpression system (Suppl. S1A) in SC-islets to model MAFA-mediated human β cell maturation. The SC-islet differentiation protocol used in these studies has been adapted from established protocols^4,21,22^ and occurs over 27 days in 6 stages (S1-6) with SC-islets exhibiting the greatest, albeit limited, glucose-stimulated insulin secretion (GSIS) response from days 26-30 (Fig. 1A).

**Figure 1.**
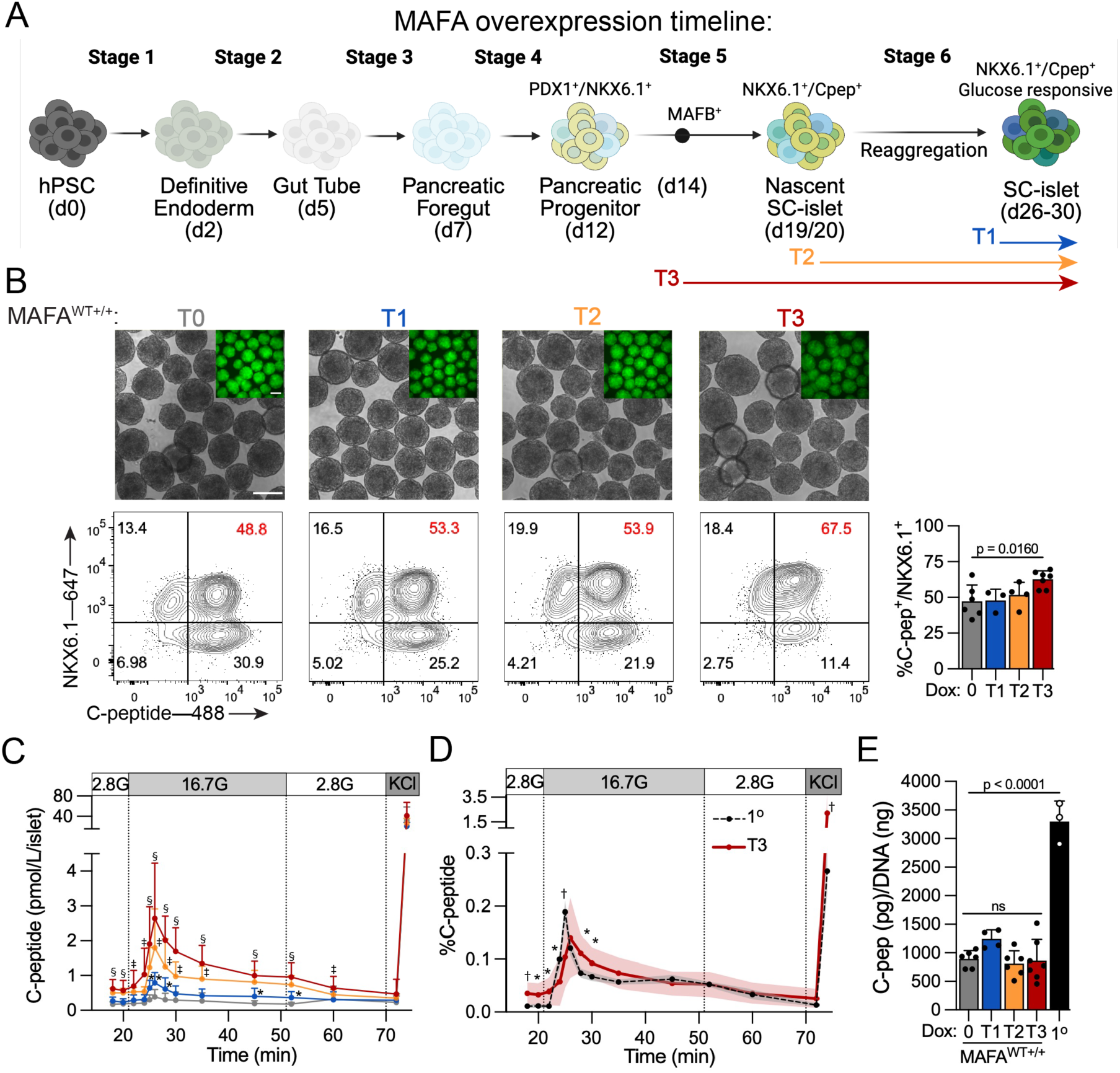
MAFA^WT+/+^ overexpression progressively increases. β **cell identity markers and GSIS functionality. (A)** Schematic showing *in vitro* SC-islet differentiation and MAFA overexpression time course. **(B)** Representative brightfield and GFP images of SC-islets overexpressing MAFA^WT+/+^ and quantitative analysis of SC-islets showing proportion of cells co-positive for Cpep^+^/NKX6.1^+^. p-values by one-way Anova followed by Dunnett’s multiple comparisons test versus non-overexpressing controls. Scale bar 150 µm. **(C)** Dynamic GSIS representing C-peptide (pmol/L) normalized per islet that is secreted in response to 2.8 mM glucose, 16.7 mM glucose, and 30 mM KCl in 2.8 mM glucose KRB. p-values by two-way repeated measures Anova followed by Dunnett ‘s multiple comparisons test versus non-overexpressing controls. **(D)** Dynamic GSIS results of MAFA^WT+/+^ T3 SC-islets normalized to total C-peptide content in comparison to primary human donor islets (n= 3 donors). p-values by two-way repeated measures Anova followed by Šídák’s multiple comparisons test. **(E)** Total C-peptide content in MAFA^WT+/+^ SC-islets and human donor islets normalized to DNA (ng). p-values by one-way Anova followed by Tukey’s multiple comparisons. Data are presented as mean ± s.e.m using at least 3 biological samples (n≥3). *p<0.05, ^†^p<0.005, ^‡^p<0.0005, ^§^p<0.0001.

MAFA^WT+/+^ overexpression was initiated at three key time points in the differentiation protocol (Fig. 1A): T1) for 72 hours at the SC-islet stage, T2) starting at day 20 in the nascent SC-islet stage when SC-β cell insulin expression is evident but the cells are unresponsive to glucose stimulation, and T3) shortly after pancreatic endoderm formation at the time when MAFB is expressed and initial insulin expression begins (MAFB expression is pre-requisite to insulin and MAFA expression during human β cell development^23^). All assays were conducted at the SC-islet stage. Initial immunofluorescence studies confirmed successful dox-induced MAFA protein expression (Suppl. S1B). Flow cytometry analysis for β cell identity markers (PDX1, NKX6.1 and C-peptide) revealed that short overexpression of MAFA^WT^ (T1) did not alter SC-islet β cell composition compared to non-overexpressing control SC-islets (Fig.1B; Suppl. S1C). MAFA^WT+/+^ T2 did not increase the amount of β cells co-expressing NKX6-1^+^/C-peptide^+^, but it did increase the amount of PDX1^+^/NKX6.1^+^ cells indicating a shift towards more β cells. Increasing the duration of MAFA^WT+/+^ by starting shortly after pancreatic endoderm formation (T3) significantly increased the total number of insulin-expressing β cells within SC-islets. Concurrently MAFA^WT+/+^ T3 SC-islets also showed a reduction in α cell glucagon hormone expression (Suppl. S1D). Additional control experiments demonstrated that dox treatment did not significantly impact the differentiation of unmodified control SC-islets (Suppl. S1E). These findings are consistent with MAFA being a transcription factor that targets and stabilizes β cell identity markers.

In addition to promoting β cell gene expression, MAFA has also been shown to improve β cell functional maturation by enhancing the amount of insulin secreted both at basal glucose concentrations and in response to high glucose challenge. To determine the impact of MAFA^WT+/+^ overexpression (OE) on SC-islet insulin secretion, we performed dynamic glucose-stimulated insulin secretion (GSIS) perifusion assays and compared SC-islets treated with dox to edited SC-islets not exposed to dox. Across all three time points (T1-3), MAFA^WT+/+^ OE SC-islets exhibited significantly improved GSIS, albeit to varying degrees (Fig. 1C). At T1, there was a marginal but significant improvement in the peak of the first phase of insulin secretion along with indications of an emerging second phase. Both T2 and T3 SC-islets showed greater basal insulin secretion, increased first phase insulin secretion and a robust second phase, with T3 over T2 SC-islets showing significant improvement. When normalizing MAFA^WT+/+^ T3 secreted C-peptide to total C-peptide content, we observed an insulin secretion pattern that is more similar to that of adult human donor islets indicative of improved maturation status and control over SC-islet insulin secretion (Fig. 1D). Total C-peptide levels did not change across the MAFA^WT+/+^ SC-islets (Fig. 1E), indicating that MAFA^WT+/+^ T3 SC-islets maintain their intracellular stores while secreting significantly more insulin. However, MAFA^WT+/+^ SC-islets still produce approximately 3-to 4-fold less insulin per cluster than adult human donor islets (Fig. 1E, Suppl. S1F). Overall, these results demonstrate that MAFA^WT+/+^ positively regulates genes involved in maintaining β cell identity and enhancing insulin secretion, consistent with findings from studies performed in rodent models.

### MAFA^S64F^ MODY variant transiently improves SC-islet functionality before rendering cells non-functional

Despite MAFA generally being associated with positive effects in β cells, a MAFA mutant variant (MAFA^S64F^) has been identified as a MODY gene. This point mutation, which substitutes serine 64 (S64) with a phenylalanine (F), prevents phosphorylation of the S^65^ priming site, inhibiting subsequent phosphoregulation and increasing MAFA protein stability^19,20^ (Fig. 2A). Heterozygous males are predominantly affected by MAFA^S64F^, typically developing diabetes in their late 30’s^19^. Mouse studies suggest that, in male mice, MAFA^S64F+/-^ leads to an initial hypersecretion of insulin that is shortly followed by premature β cell aging and senescence^20^. To further delineate how human β cell biology is dysregulated by the MAFA^S64F^ TF in comparison to MAFA^WT^ TF, we modeled the effects of MAFA^S64F+/-^ in SC-islets over the same overexpression time course as MAFA^WT+/+^ (Fig. 1A). Immunoblotting for MAFA protein confirms that MAFA^S64F+/-^ is expressed at much higher levels than MAFA^WT+/+^ ^19^, including when comparing MAFA^WT+/+^ T3 to MAFA^S64F+/-^ T1 SC-islets (Fig. 2B). In stark contrast to MAFA^WT+/+^ SC-islets, MAFA^S64F+/-^ SC-islets exhibited a significant reduction in C-peptide^+^ cells even as early as the T1 overexpression time point, with this effect being further exacerbated with extended overexpression at T2 and T3 (Fig. 2C and Supp. S2A). However, the expression of PDX^+^/NKX6.1^+^ was maintained except with the longer T3 overexpression time point when there was significant loss of NKX6.1^+^ (Supp. S2A). These findings show that MAFA^S64F+/-^ rapidly depletes intracellular insulin and with longer overexpression, causes a loss of NKX6.1 suggesting a loss of β cell identity.

**Figure 2.**
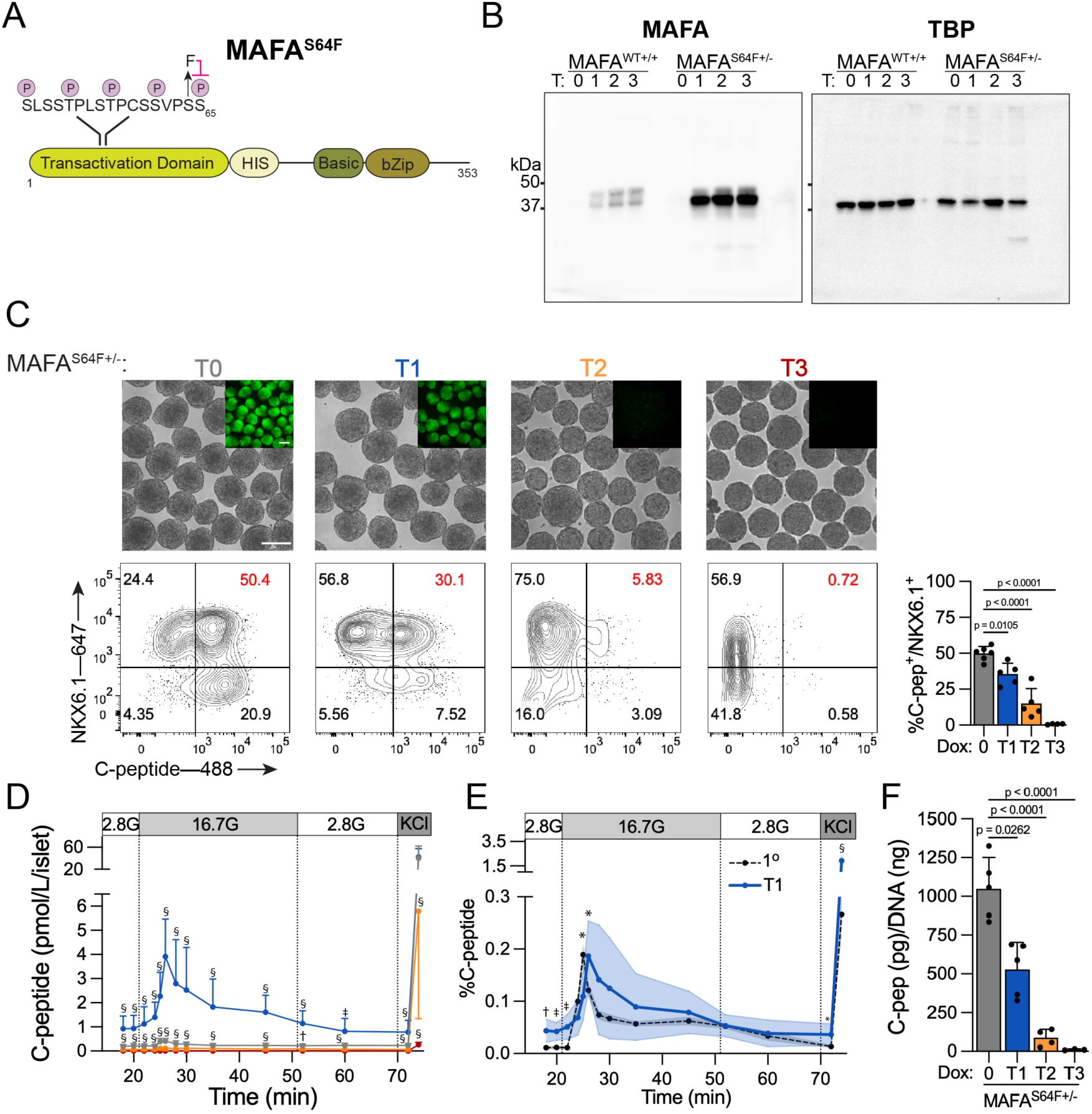
MAFA^S64F+/-^ transiently increases SC-islet functionality before rendering β cells non-functional. **(A)** MAFA schematic showing the missense mutation of Serine (S) 64 to Phenylalanine (F) in the transactivation domain. **(B)** Western blot of MAFA^WT+/+^ and MAFA^S64F+/-^ SC-islets cell lysates for MAFA (left) and TBP loading control (right) protein expression. **(C)** Representative brightfield and GFP images of SC-islets overexpressing MAFA^S64F+/-^ and quantitative analysis of SC-islets showing proportion of cells co-positive for Cpep^+^/NKX6.1^+^. p-values by one-way Anova followed by Dunnett’s multiple comparisons test versus non-overexpressing controls. Scale bar 150μm. **(D)** Dynamic GSIS perifusion assay representing C-peptide (pmol/L) normalized per islet that is secreted in response to 2.8mM glucose, 16.7mM glucose, and 30mM KCl in 2.8mM glucose KRB. p-values by two-way repeated measures Anova followed by Dunnett ‘s multiple comparisons test versus non-overexpressing controls. **(E)** Dynamic GSIS results of MAFA^S64F+/-^ T1 SC-islets normalized to total C-peptide content in comparison to primary human donor islets (n= 3 donors). p-values by two-way repeated measures Anova followed by Šídák’s multiple comparisons test. **(F)** Total C-peptide content in MAFA^WT+/+^ SC-islets and human donor islets normalized to DNA (ng). p values by one-way Anova followed by Tukey’s multiple comparisons. *p<0.05, ^†^p<0.005, ^‡^p<0.0005, ^§^p<0.0001. Data are presented as mean ± s.e.m. n= at least 3 biological samples.

Next, we examined how MAFA^S64F+/-^ modulates SC-islet functionality. Short overexpression of MAFA^S64F+/-^ in SC-islets (T1) led to a robust increase in insulin secreting abilities during basal insulin secretion and in response to glucose stimulation in dynamic GSIS (Fig. 2D). The extent of C-peptide secreted by MAFA^S64F+/-^ T1 SC-islets was similar to that secreted by MAFA^WT+/+^ T3 SC-islets (Supp. S2B). However, T2 and T3 MAFA^S64F+/-^ SC-islets secreted significantly lower levels of C-peptide that are almost undetectable except in response to KCl. Similar, to MAFA^WT+/+^ T3 SC-islets, the percentage of C-peptide secreted by MAFA^S64F+/-^ T1 SC-islets is more comparable to that observed in primary human donor islets than control SC-islets (Fig. 2E). However, when comparing total C-peptide content in control, non-overexpressing SC-islets to MAFA^S64F+/-^ SC-islets we observed a significant loss at T1 that continued to decrease at T2 and T3 (Fig. 2F), indicating that MAFA^S64F+/-^ SC-islets are unable to maintain intracellular insulin stores. These findings demonstrate that MAFA^S64F+/-^ SC-islets show a transient increase in insulin secretory ability before depleting insulin stores and rendering the cells non-functional, recapitulating many of the phenotypes identified in human patients and mouse models^19,20^.

### MAFA^WT+/+^ remodels the SC-islet metabolic landscape

Immature SC-islet functionality and a divergent transcriptome compared to human islets have, in part, been attributed to low levels of MAFA transcriptional activity^3–5,12^. To better understand the influence of MAFA on SC-islet gene expression, we performed bulk RNAseq and proteomics analysis of SC-islets. Comparing the longest MAFA^WT+/+^ overexpression T3 timepoint to no dox samples as a reference, we identified 1717 genes and 695 proteins that had at least a 0.325 log2 fold change (25%) increase and were statistically significant (padj≤0.05) (Fig. 3A). Gene Ontology (GO) analysis identified an upregulation of biological processes pertaining to organelle organization and cellular metabolism including cilium organization, Golgi organization, fatty acid metabolic process and electron transport chain (ETC) (Fig. 3B). Employing Kyoto Encyclopedia of Genes and Genomes (KEGG) analysis to further understand the pathways underlying the biological processes influenced by MAFA revealed increases in carbon metabolism, citrate cycle (TCA cycle), and oxidative phosphorylation (OxPhos), as well as unanticipated pathways such as fatty acid metabolism and peroxisomes. Notably, to determine processes important for sustaining greater SC-islet functionality, input genes and proteins were further narrowed to only include those that were significantly upregulated in MAFA^WT+/+^ T3 but not in MAFA^S64F+/-^ T1 SC-islets (426 genes and 329 proteins). GO biological processes that persist include cilium organization and ETC while persistent KEGG pathways include OxPhos, fatty acid metabolism and peroxisomes (Supp. S3A). Moreover, pathway analysis on MAFA^S64F+/-^ T1 SC-islet upregulated RNA and proteins indicated an increase of cellular stress such as the GO term response to oxidative stress and of autophagy and lysosome KEGG pathways (Suppl. S3B). These findings highlight discrepancies in metabolic remodeling and organelle organization as potential important factors contributing to sustained MAFA^WT+/+^ function and MAFA^S64F+/-^ dysfunction.

**Figure 3.**
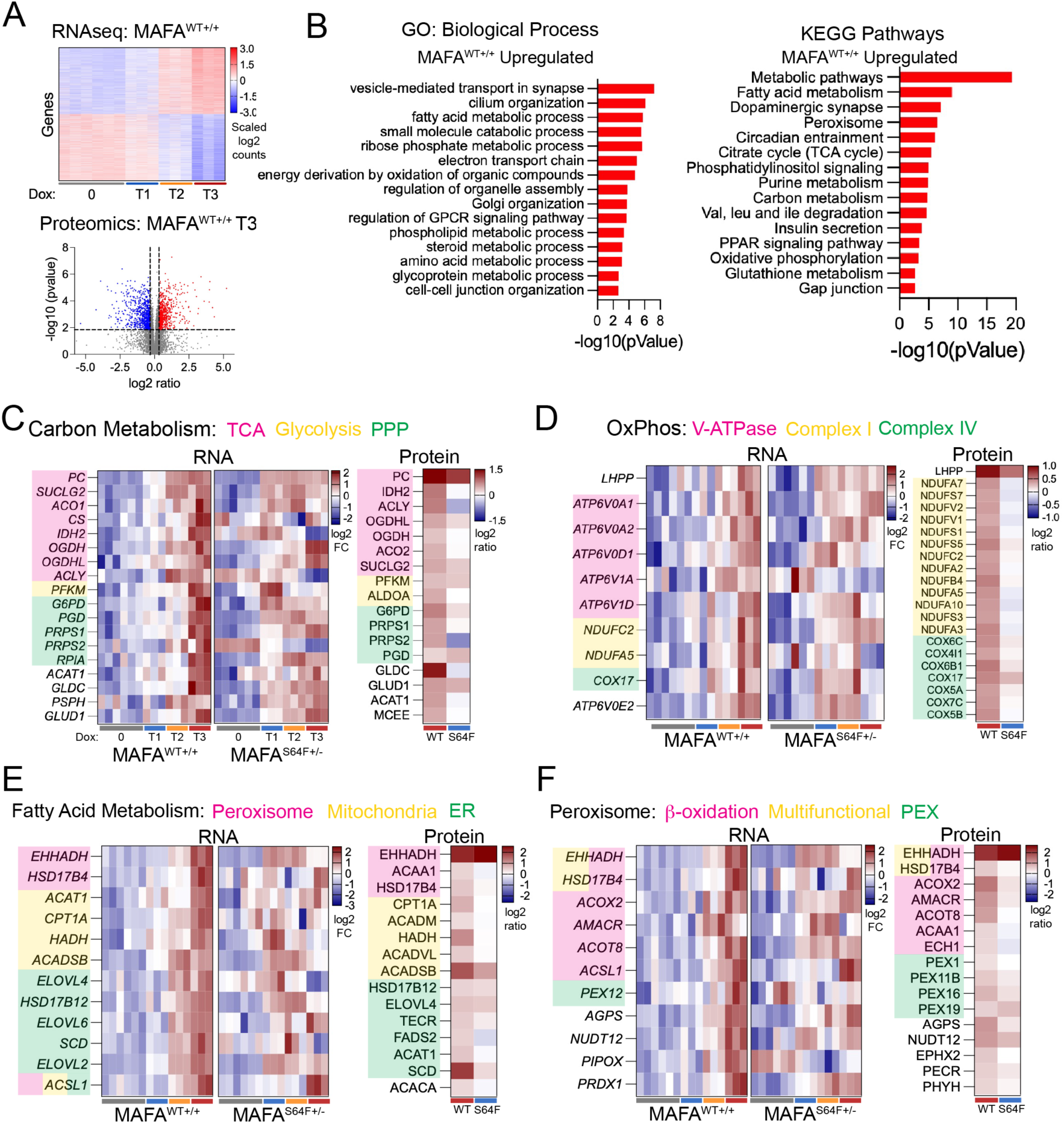
MAFA^WT+/+^ upregulates SC-islet metabolic processes. **(A)** MAFA^WT+/+^ T3 SC-islet RNA heatmap (top) and protein volcano plot (bottom) showing RNA and proteins that are significantly upregulated (red) and downregulated (blue). **(B)** GO and KEGG pathway analysis of RNA and protein significantly upregulated in MAFA^WT+/+^ T3 SC-islets (FDR<0.05). **(C-F)** Heatmaps of RNA and protein from select KEGG pathways identified in (B). C) Heatmap of genes and proteins involved in carbon metabolism and further annotated for specific roles in TCA, glycolysis and PPP. D) OxPhos pathway annotated for V-ATPase, and ETC Complex I and IV. E) Fatty acid metabolism annotated for cellular localization in peroxisome, mitochondria, or ER. F) Peroxisome annotated for β-oxidation, multifunctional, or peroxin (PEX) proteins. RNA is showed in a log2 fold change (FC) format and protein is represented as log2 ratio of MAFA^WT+/+^ T3 or MAFA^S64F+/-^ T1 over non-overexpressing SC-islets.

Expression heatmaps of factors involved in the selected KEGG pathways (Carbon metabolism, OxPhos, Fatty Acid Metabolism, and Peroxisomes) provide deeper insights into the relationship between MAFA overexpression, subsequent transcriptional and translational changes, and GSIS functionality (Fig. 3C-F). In MAFA^WT+/+^, longer MAFA overexpression correlated with a progressive upregulation of gene products in these metabolism-associated KEGG pathways and greater insulin secretory capacity. In contrast, MAFA^S64F+/-^ T1 SC-islets showed irregular RNA upregulation across these pathways with limited changes in protein expression despite having comparable insulin secretory capacity to MAFA^WT+/+^ T3 SC-islets. Notably, two genes did show robust increases in protein expression - the TCA enzyme pyruvate carboxylase (PC) and the OxPhos protein LHPP, both of which have been implicated in increasing mitochondrial OxPhos by aiding in TCA flux^24–26^. Therefore, these results suggest that increases in TCA flux allows for greater glucose metabolism via glycolysis, the TCA cycle and OxPhos as a means to enhance GSIS functionality in both MAFA^WT+/+^ T3 and MAFA^S64F+/-^ T1 SC-islets.

In addition to the canonical carbon metabolism and OxPhos pathways implicated in GSIS, we also see increased expression of genes involved in biogenesis and energy production in MAFA^WT+/+^ SC-islets (Fig. 3C-F). These include nucleoside generation (PRPS1, PRPS2, RPIA) by the pentose phosphate pathway (PPP) (Fig. 3C), fatty acid elongation in the ER (ELOVL4, HSD17B12, ELOVL6, ELOVL2) (Fig. 3E) and fatty acid β-oxidation in peroxisomes (EHHADH, HSD17B4, ACOX2, AMACR, ACOT8, ACSL1) and mitochondria (CPT1A, ACADM, HADH, ACADVL, ACADSB, ACSL1) (Fig. 3F). However, these genes were not uniformly upregulated in MAFA^S64F+/-^ T1 SC-islets. One exception was the peroxisome protein EHHADH, which was robustly upregulated in both MAFA^WT+/+^ T3 and MAFA^S64F+/-^ T1 SC-islets. In addition to its role in β-oxidation, EHHADH has also been shown to reduce reactive oxygen species (ROS)-mediated pexophagy^27^ , the break down and removal process of peroxisomes, and managing ROS produced by mitochondria during increased metabolic activity^28^. Although EHHADH was upregulated in MAFA^S64F+/-^ T1 SC-islets there was not the same increased expression of peroxisome genes (RNA and protein) as was observed in MAFA^WT+/+^ T3 SC-islets (Fig. 3F). V-ATPase, another important component for maintaining intracellular redox homeostasis, showed a similar pattern of being upregulated in MAFA^WT+/+^ T3 SC-islets but not MAFA^S64F+/-^ T1 SC-islets (Fig. 3D, Supp. S3C). Collectively, these findings suggest that sustaining heightened GSIS functionality in MAFA^WT+/+^ T3 SC-islets requires concomitant increases in biogenesis, energy production, redox homeostasis, and secretory metabolic stimulatory pathways and that the absence of the gene products regulating these processes contributes to MAFA^S64F+/-^ SC-islet dysfunction.

### MAFA serves as a signaling hub to initiate SC-islet maturation

To better understand how MAFA influences gene regulatory networks (GRNs) that govern the above biological processes and pathways, we generated single nuclei multi-omics data (snRNAseq and snATACseq) and analyzed it using SCENIC+^29^. Additionally, we leveraged tools from the scverse core and ecosystem, including Scanpy, MultiVI, and DRVI, to enhance data integration and analysis^30^. We resolved 15 cell type clusters and identified specific islet cell types including SC-β (β^1^ and β^2^), SC-α, SC-δ and serotonergic islet cells (SIC^1^ and SIC^2^)^31^ based on gene expression (Fig. 4A). MAFA expression and sample type were used to further identify clusters corresponding to MAFA overexpression timepoints (Suppl. S4A) as well as transitional MAFA overexpressing cell-types SC-α-to-β and SC-δ-to-β. Interestingly, MAFA^WT+/+^ overexpressing SC-β clusters showed increased markers for serotonin synthesis machinery (*TPH1*, *DDC*), but in contrast to SIC clusters, they retained high expression of β cell identity markers (*INS*, *PDX1*, *NKX6-1*, *EBF1*) and prominently expressed the β cell-associated *SLC18A2* monoamine transporter over the enterochromaffin-associated *SLC18A1* monoamine transporter^32–37^ (Fig. 4B, Suppl. S4B).

**Figure 4.**
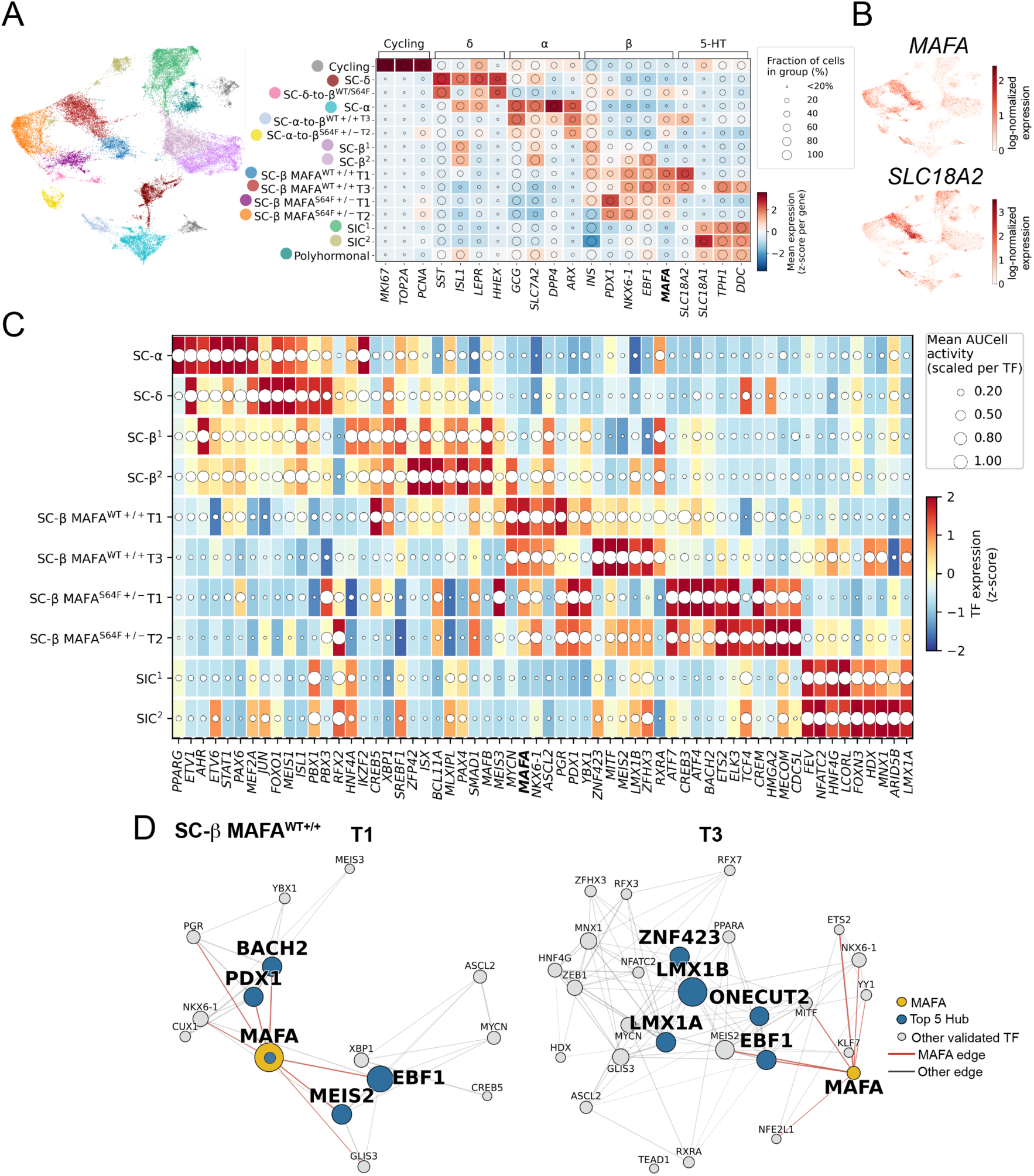
Transcription factor network architecture across SC-islet cell states, MAFA genotypes, and differentiation timepoints. **(A)** Joint multi-omic (RNA + ATAC) UMAP of SC-islets, colored by annotated cluster (left). Right: RNA expression of cell-identity markers across clusters, grouped by lineage. Tile color = mean expression (z-score per gene); overlaid circle size = fraction of cells in the cluster expressing the gene. Superscripts denote MAFA genotype (^WT+/+, S64F+/−^) and differentiation timepoint (T1–T3). **(B)** Same UMAP as in A, colored by log-normalized expression of MAFA and SLC18A2. Scales are independent per gene. **(C)** Union of the top 10 most active TFs per cluster, excluding cycling, transitional, and polyhormonal clusters. Tile color = TF expression (z-score across clusters); overlaid circle size = mean AUCell regulon activity (min–max scaled per TF). Note the two encodings are independent: a large circle on a blue tile indicates high regulon activity despite low TF expression. **(D)** Hub signaling network plots of TF co-regulatory networks determined by target gene overlap in MAFA^WT+/+^ T1 and T3 clusters. Nodes represent validated TFs derived from eRegulon analysis using SCENIC+. Node size is proportional to a composite centrality score, calculated as the min-max normalized mean of degree, betweenness, and eigenvector centrality. Edges represent significant target-gene overlaps between TF pairs (Benjamini-Hochberg FDR < 0.05).

SCENIC+ eRegulon inference allowed for the identification of active transcription factors (TFs) within each cluster. Analysis of the top 10 active TFs revealed that SC-α, SC-δ, and SC-β^1^/-β^2^ show large overlap for endocrine progenitor-associated TFs including (*ETV1*, *JUN*, *FOXO1*, *HNF4A*, *ZFP42*, *BCL11A*, *PAX4*) consistent with SC-islets having a higher degree of plasticity and not being terminally differentiated (Fig. 4C)^2,38^.

Analysis of active TFs in SC-β MAFA^WT+/+^ T1 clusters revealed that even a short overexpression of MAFA^WT+/+^ (3 days) begins to reinforce the β cell identity as evidenced by decreased pancreatic endocrine progenitor TF activity and increased *NKX6.1* and *PDX1* activity (Fig. 4C). With longer overexpression, additional β cell-associated TFs were activated including *MEIS2*, *MITF*, *LMX1B*, *PGR* and *MNX1* as well as TFs involved in circadian entrainment (*ZFHX3*) and serotonin biosynthesis (*FEV*, *LMX1A*)^38–47^. In contrast, SC-β MAFA^S64F+/-^ T1 exhibit a sharp activation of stress-associated markers including ER stress TFs (*ATF4*^48^ and *CREB3*^49^), oxidative stress sensors (*BACH2*^50^ and *ETS2*^51^), the stress sensor *YBX1*^52,53^ and the stress-responsive chromatin regulator *ATF7*^54,55^, as well as some TFs that have been linked to stress-induced plasticity or stem-like states (*HMGA2*^56^ and *MECOM*^57^). Notably, stress-related TF activity is relatively low in SC-β MAFA^WT+/+^ T1 and further resolves by MAFA^WT+/+^ T3, suggesting a protective or adaptive affect in MAFA^WT+/+^ cells, whereas high-level of activation of these factors in MAFA^S64F+/-^ clusters is detrimental.

To further explore each cluster transcriptional program, we constructed TF-TF networks based on shared regulatory target genes and applied network centrality analysis. This identified TF signaling hubs that may act as coordination points within that cluster’s regulatory architecture. The top 5 hubs in SC-β MAFA^WT+/+^ T1 were MAFA, EBF1, MEIS2, BACH2, and PDX1 with MAFA being the leading, most interconnected hub consistent with induced MAFA^WT+/+^ expression driving phenotypic changes (Fig. 4D). In SC-β MAFA^WT+/+^ T3 cells, the top 5 signaling hub hierarchy had been restructured to no longer include MAFA which had been replaced by LMX1B as the primary hub followed by EBF1, LMX1A, ZNF423 and ONECUT2. In contrast, SC-β MAFA^S64F+/-^ T1 and T2 clusters show very distinct signaling hub patterns from MAFA^WT+/+^ clusters (Suppl. S4C). Most notably, in MAFA^S64F+/-^ T1 clusters, MAFA is not the primary signaling hub and four of the top 5 signaling hubs are stress-associated TFs (CREM, BACH2, DRAP1, ATF7)^50,58,59^. These findings indicate that MAFA^WT+/+^ drives GRN remodeling that function to reinforces β cell identity, whereas phenotypic changes in MAFA^S64F+/-^ clusters may be driven by overtly activated cell stress response.

### SC-islet maturation is driven by coordinated transcription factor activity

To better understand how active TFs elicit their effects within the MAFA^WT+/+^ T1 and T3 clusters, we performed Louvain clustering analysis and identified TF communities based on shared target genes. MAFA^WT+/+^ T1 and T3 TFs readily clustered into communities indicating that TFs within these clusters function in a coordinated manner (Fig. 5A). In contrast, Louvain clustering of MAFA^S64F+/-^ T1 cluster did not readily identify similarly coordinated communities, and many of the active TFs remained isolated (Suppl. S5A). Thus, coordinated TF activity distinguishes MAFA^WT+/+^ clusters from discordant TFs in MAFA^S64F+/-^ T1 cluster.

**Figure 5.**
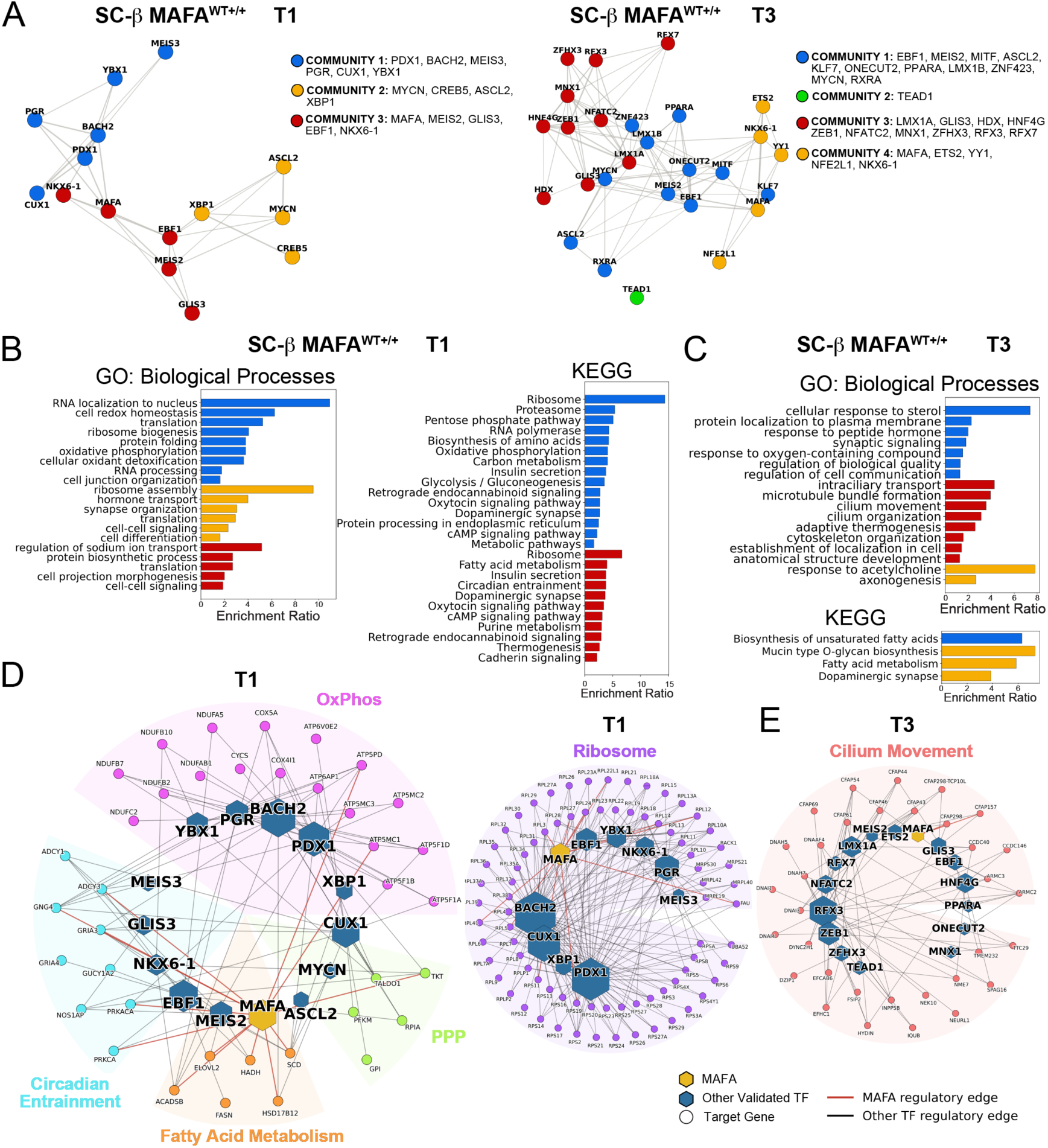
Transcription factor communities coordinate SC-islet maturation process. **(A)** Louvain community clustering of active transcription factors in MAFA^WT+/+^ T1 (resolution 1.32) and T3 (resolution 1.02) clusters. **(B)** MAFA^WT+/+^ T1 Louvain communities DEG pathway analysis. **(C)** MAFA^WT+/+^ T3 Louvain communities DEG pathway analysis. **(D)** MAFA^WT+/+^ T1 TF-to-gene network plots for OxPhos, PPP, Fatty Acid Metabolism, Circadiant Entrainment, and Ribosome upregulated KEGG pathways. **(E)** TF-to-gene network plots for Cilium Movement upregulated KEGG pathway in MAFA^WT+/+^ T3.

Community-shared DEG target genes were analyzed for GO biological processes and KEGG pathway enrichment, similar to previous bulk RNAseq and proteomics analyses. The enrichment analysis used the eRegulon target universe as the background rather than the entire genome, as community definitions are based on shared targets, and coherent enrichment is inherently integrated into the community framework. MAFA^WT+/+^ T1 community DEG pathway analysis uncovers biological processes associated with cellular growth and homeostasis including translation, ribosome biogenesis and cell redox homeostasis, as well as KEGG pathways that were previously identified in bulk RNAseq and proteomics analysis including OxPhos, PPP, glycolysis, fatty acid metabolism and circadian entrainment (Fig. 5B, Suppl. S5B). Intriguingly, MAFA^WT+/+^ T3 pathway analysis revealed that community 3 (*LMX1A*, *GLIS3*, *HDX*, *HNF4G*, *ZEB1*, *NFATC2*, *MNX1*, *ZFHX3*, *RFX3*, *RFX7*) strongly regulated cilium organization and movement—a process that has been implicated in β cell glucose-responsiveness in human and rodents (Fig. 5C)^60–62^. Visualizing inferred TF-to-gene connections provides further insight into how GRNs may be structured to give rise to key processes such as OxPhos, Circadian Entrainment, Fatty Acid Metabolism, PPP, Ribosome biogenesis and ciliary movement during β cell maturation (Fig. 5D, E). Collectively, these studies further support the bulk RNAseq and proteomics findings while providing new information on how GRNs may govern cellular biogenesis, metabolism, and redox homeostasis during MAFA-mediated SC-islet functional maturation.

### MAFA^WT+/+^ SC-islets prioritize cellular fuels for biogenesis before switching to oxidative phosphorylation

The strong omics footprint for cellular biogenesis and modified SC-islet metabolism in response to MAFA OE prompted us to utilize <u>m</u>ulti-isotope <u>m</u>ass <u>s</u>pectrometry correlated with <u>e</u>lectron <u>m</u>icroscopy (MIMS-EM)^63–65^ to simultaneously examine how cellular architecture and nutrient metabolism are modified by MAFA^WT+/+^ and MAFA^S64F+/-^ *in situ* (Fig. 6A, B). SC-islets were incubated in media containing [U-^13^C6]-glucose (^13^C-glucose, 11mM) and ^15^N-glutamine (2mM) prior to fixation and sample processing. Quantitative analysis of EM images revealed that MAFA OE leads to dynamic changes in mitochondria size (Fig. 6C). MAFA^WT+/+^ induced a significant increase in mitochondria size at timepoints T1 and T2, but a decrease in mitochondria size at T3. In striking contrast, mitochondria size in MAFA^S64F+/-^ SC-islets significantly increases, giving rise to swollen mitochondria at T2 (+21±1.2%). These findings suggest that MAFA^WT+/+^ cells dynamically maintain mitochondrial homeostasis during increased metabolic demands whereas mitochondria swelling in MAFA^S64F+/-^ cells is an indicator of cellular stress and mitochondrial dysfunction that is associated with cellular senescence^66^.

**Figure 6.**
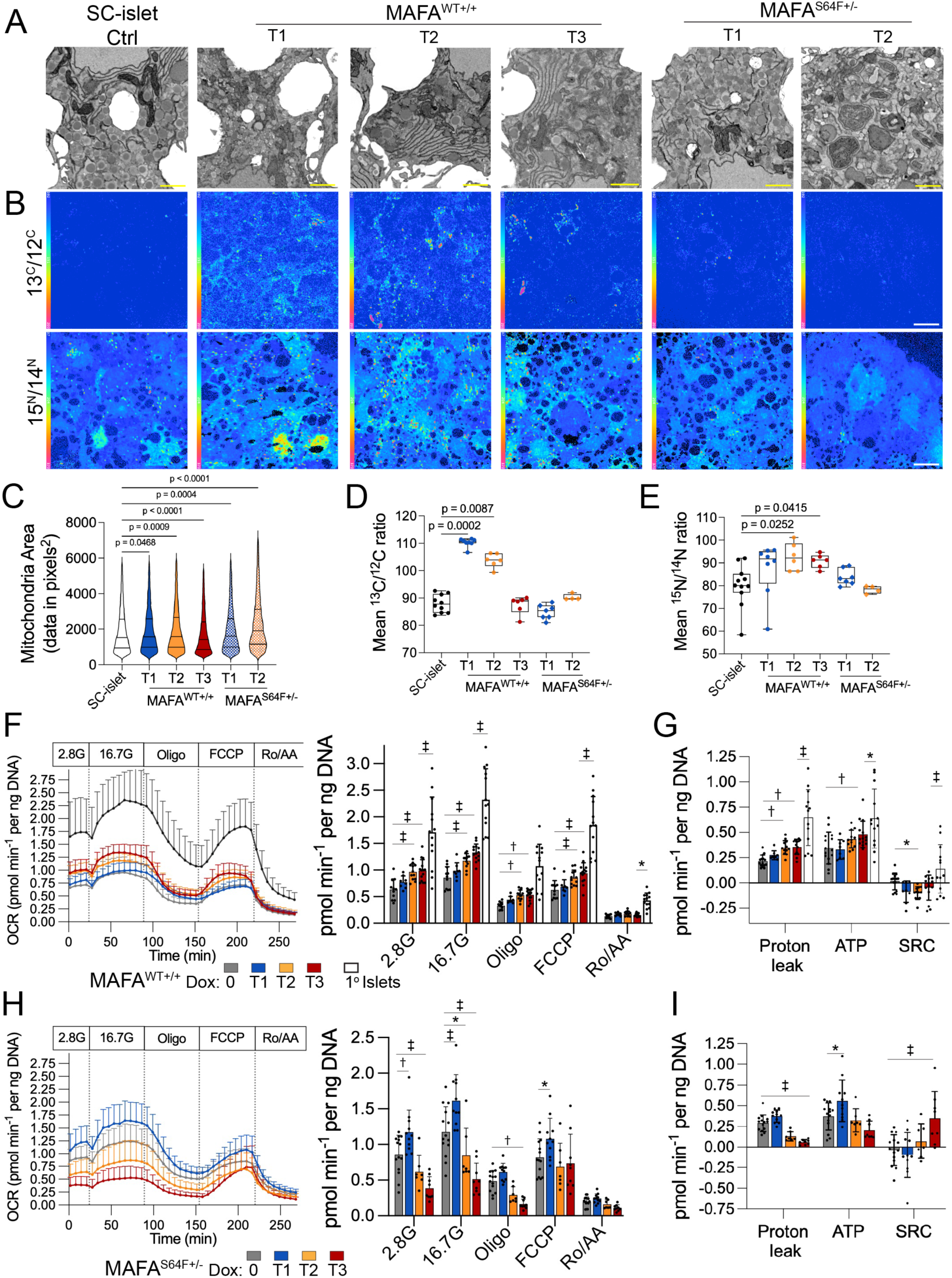
MAFA^WT+/+^ drives cellular biogenesis to expand respiratory capacity. **(A)** EM visualization of SC-islets. Scale bar 1μm. **(B)** MIMS images of SC-islets labeled with 13^U^-carbon glucose (11mM) or 15^N^-nitrogen glutamine (2mM). Middle panels: 13^C^/12^C^ carbon isotope label. Bottom panels: 15^N^/14^N^ glutamine isotope label. Scale bar 1μm. **(C)** Quantification for number of mitochondria identified in EM images. **(D)** Mean 13^C^/12^C^ signal across samples. **(E)** Mean 15^N^/14^N^ signal across samples. **(F)** Seahorse assay of MAFA^WT+/+^ SC-islets and human donor islets showing traces of oxygen consumption rates normalized to DNA as a readout for mitochondrial respiration in response to baseline low (2.8mM) glucose KRB, high (16.7 mM) glucose, oligo (5 μM), FCCP (1 μM), or rotenone (5 μM) and antimycin A (5 μM). Quantification of traces across the baseline and for the maximal response to each treatment. p-values were determined by two-way Anova followed by Dunnett’s multiple comparisons versus non-overexpressing SC-islets or human donor islets. **(G)** Analysis of OCR traces for proton leak, ATP production, or spare respiratory capacity. p-values by two-way Anova followed by Dunnett’s multiple comparisons test versus non-overexpressing controls. **(H)** Normalized OCR traces for MAFA^S64F+/-^ SC-islets and quantified for baseline and maximal responses to indicated treatment. p-values by two-way Anova followed by Dunnett’s multiple comparisons test versus non-overexpressing controls. **(I)** Analysis of OCR traces for proton leak, ATP production, or spare respiratory capacity. p-values by two-way Anova followed by Dunnett’s multiple comparisons test versus non-overexpressing controls. Data are presented as mean ± s.e.m using at least 3 biological samples (n≥3) unless otherwise stated.

To investigate how MAFA changes fuel usage in SC-islets, we analyzed spatial isotope labeling patterns and their subcellular compartmentation. In MIMS-EM, increased accumulation of ^13^C and ^15^N in cellular biomass is an indication that glucose and glutamine, respectively, are being utilized by the cells for biomass synthesis. Using this approach, we found that glucose-derived ^13^C labeling was relatively low in control SC-islets and was significantly elevated in MAFA^WT+/+^ T1 and T2 (Fig. 6B, D, E). In contrast, MAFA^S64F+/-^ T1 and T2 SC-islets showed no detectable increases in ^13^C levels. Additionally, ^15^N labeling was significantly elevated in MAFA^WT+/+^ cells at T2 and T3 timepoints, whereas no changes were detected in MAFA^S64F+/-^ SC-islets. Taken together, this data suggests that MAFA^WT+/+^ cells can utilize and retain glucose- and glutamine-derived elements for biomass synthesis whereas, these aspects would be impaired in MAFA^S64F+/-^ SC-islets. Moreover, MAFA^WT+/+^ T3 SC-islets’ enhanced GSIS response alongside the upregulation of metabolic and OxPhos genes (Fig. 3B-D and Fig. 5D) suggested that their reduced accumulation of glucose-derived ^13^C levels may be due to increased glucose turnover. To test this hypothesis, we used Seahorse to measure oxygen consumption rates (OCR) as a readout for mitochondrial ETC activity. We observed an overall increase in mitochondrial respiration in MAFA^WT+/+^ T3 SC-islets, with a more modest increase at T2 (Fig. 6F). It is worth noting that MAFA^WT+/+^ T3 mitochondrial activity did not reach the same magnitude as primary human donor islets. Further analysis of Seahorse data revealed that MAFA^WT+/+^ T2 and T3 SC-islets display significantly increased proton leak and ATP production that are consistent with greater mitochondrial activity (Fig. 6G). However, the low level of spare respiratory capacity (SRC) in SC-islets regardless of MAFA OE suggest a fragile metabolic state wherein mitochondria are operating at their maximal capacity with limited flexibility to increase ATP production. We also monitored mitochondrial activity in MAFA^S64F+/-^ SC-islets and found that MAFA^S64F+/-^ T1 SC-islets had increased mitochondrial respiration and ATP production that resembled that of MAFA^WT+/+^ T3 SC-islets (Fig. 6H). However, T2 and T3 MAFA^S64F+/-^ SC-islets displayed signs of mitochondrial dysfunction, including an overall decrease in OCR, significantly reduced proton leak and a sudden increase in T3 SRC (Fig. 6I). We reasoned that increased OCR in both MAFA^WT+/+^ T3 and MAFA^S64F+/-^ T1 SC-islets may be the result of increased pyruvate carboxylase activity (Fig. 3C), which would improve coupling of glycolysis to mitochondrial TCA and ETC activity. Metabolic tracing experiments confirmed increased flux via PC in MAFA^WT+/+^ T3 and MAFA^S64F+/-^ T1 SC-islets confirming this hypothesis (Suppl. S6A). Collectively, these findings demonstrate the distinct impact of MAFA^WT+/+^ and MAFA^S64F+/-^ on fuel usage and SC-islet function.

### MAFA^WT+/+^ enhances SC-islet resilience to oxidative stress

Evaluation of MAFA ^WT+/+^ downregulated pathways gave indication that there was improvement in intracellular homeostasis and reduced internal stress (Fig. 7A). Oxidative stress emerged as a key area of interest, as reactive oxygen species (ROS) production is linked to mitochondrial metabolism and response to oxidative stress was upregulated in MAFA^S64F+/-^ SC-islets (Suppl. S3B). Thus, the ability of MAFA^WT+/+^ to better maintain redox homeostasis may contribute to MAFA^WT+/+^ SC-islets sustained functionality during heightened metabolism whereas, MAFA^S64F+/-^ undergo mitochondrial dysfunction.

**Figure 7.**
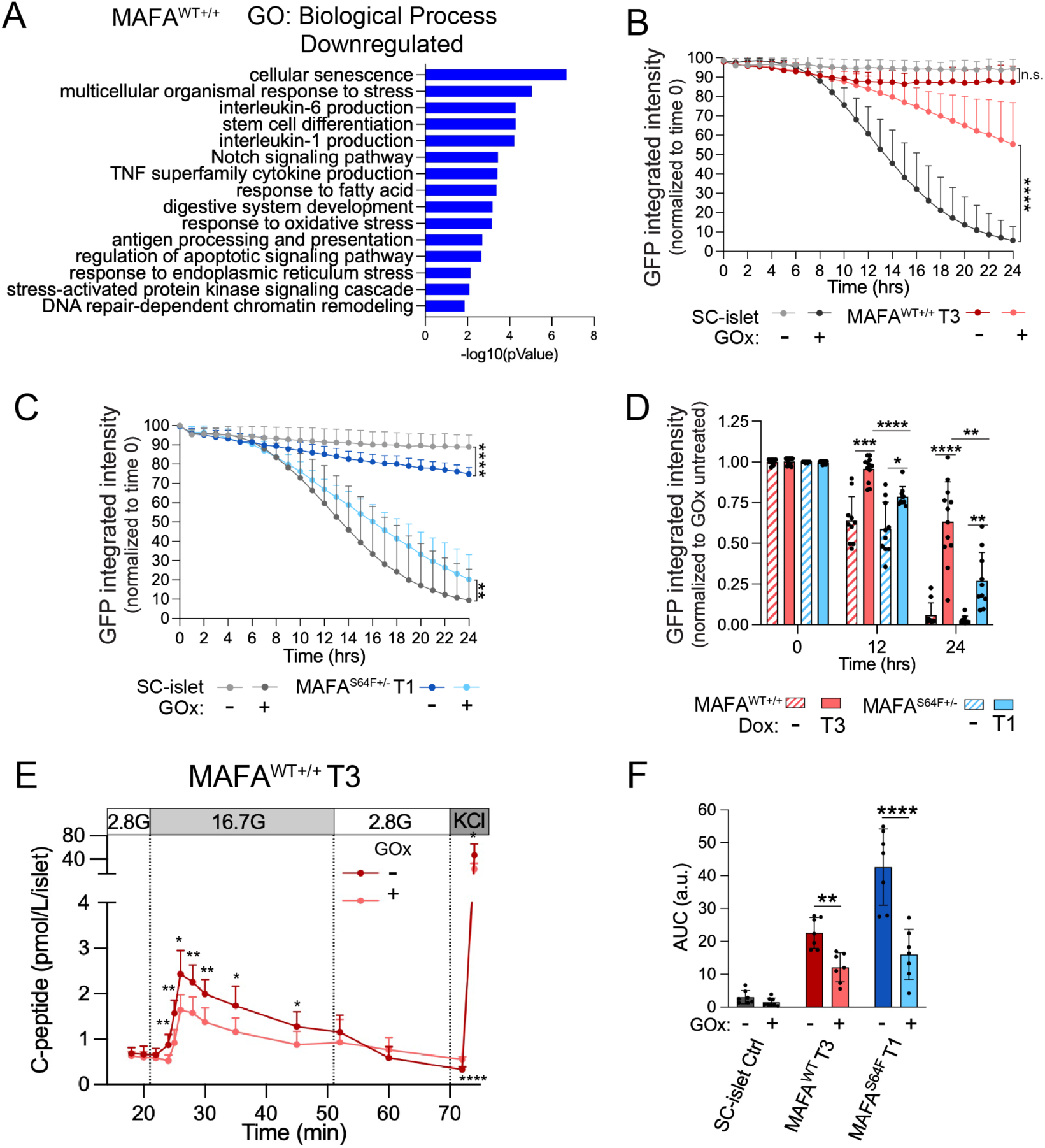
MAFA^WT+/+^ enhances SC-islet redox homeostasis. **(A)** Gene ontology biological processes downregulated in MAFA^WT+/+^ T3 SC-islets showing downregulation of stress-related pathways. p-value<0.05, FDR<0.05. **(B)** Integrated GFP intensity traces of SC-islets and MAFA^WT+/+^ T3 SC-islets treated with glucose oxidase (10 mU/ml). p-values by two-way repeated measures Anova followed by Tukey’s multiple comparisons test. **(C)** SC-islet and MAFA^S64F+/-^ T1 SC-islets incucyte GFP traces in the presence of absence of GOx (10 mU/ml). p-values by two-way repeated measures Anova followed by Tukey’s multiple comparisons test. **(D)** Quantification of non-overexpressing SC-islets, MAFA^WT+/+^ T3 or MAFA^S64F+/-^ T1 SC-islets treated with glucose oxidase (10 mU/ml). p-values by two-way repeated measures Anova followed by Tukey’s multiple comparisons test. **(E)** MAFA^WT+/+^ treated with GOx (10 mU/ml) to induce sustained oxidative stress and assessed for GSIS. p-values by two-way repeated measures Anova followed by Šídák’s multiple comparisons test. p-values by multiple unpaired t-tests, with the false discovery rate controlled using the two-stage step-up method of Benjamini, Krieger, and Yekutieli (q < 0.01). **(F)** Quantification of SC-islet controls, MAFA^WT+/+^ T3 and MAFA^S64F+/-^ T1 GSIS 16.7 mM glucose responses for area under the curve (AUC). p-values by two-way repeated measures Anova followed by Šídák’s multiple comparisons test. *p<0.05, **p<0.005, ***p<0.0005, ****p<0.00005. Data are presented as mean ± s.e.m (n≥3 biological replicates).

To determine the impact of MAFA overexpression on the response to oxidative stress, we treated cells with glucose oxidase (GOx) to induce persistent H2O2 exposure and monitored GFP as a readout for survival over a 24-hour time course. MAFA^WT+/+^ T3 islets demonstrated significantly improved retention of GFP signal compared to non-overexpressing controls, suggesting improved survival (Fig. 7B). As for MAFA^S64F+/-^ T1 SC-islets, although they performed better than non-overexpressing controls, they had significantly poorer survival rates compared to MAFA^WT+/+^ T3 SC-islets (Fig. 7C-D). Follow-up experiments demonstrated that MAFA^WT+/+^ T3 SC-islets better retained their functionality (Fig. 7E, F) compared to SC-islet controls (Suppl. S7A) and MAFA^S64F+/-^ T1 SC-islets (Suppl. S7B-C). These findings suggest that improved redox homeostasis is beneficial for sustaining MAFA^WT+/+^ SC-islet heightened metabolic activity and functionality.

### MAFA^WT+/+^ improves SC-islet functional resilience *in vivo*

To determine whether MAFA^WT+/+^ T3 SC-islets improved *in* vitro function and stress-handling properties translates to more desirable post-transplantation outcomes, MAFA^WT+/+^ T3 SC-islets were transplanted under the kidney capsule of immunodeficient mice and plasma C-peptide levels were measured during *in vivo* GSIS (Fig. 8A). MAFA^WT+/+^ T3 SC-islets demonstrated enhanced glucose responsiveness as early as 3 days post-transplantation, which persisted over the subsequent 4 weeks (Fig. 8B). In contrast, control SC-islets that did not receive doxycycline exhibited intermittent increases in C-peptide secretion, with significant responses at 7 and 14 days post-transplantation, but inconsistent results at 3 days and 4 weeks (Fig. 8B). Additionally, mice grafted with MAFA^WT+/+^ T3 SC-islets showed a trend toward higher overall C-peptide levels post-glucose injection when compared to control mice, with significant increases observed at both 14 days and 4 weeks post-transplantation. Collectively, these results suggest that the improved resilience and functionality of MAFA-matured SC-islets is beneficial for *in vivo* transplantation.

**Figure 8.**
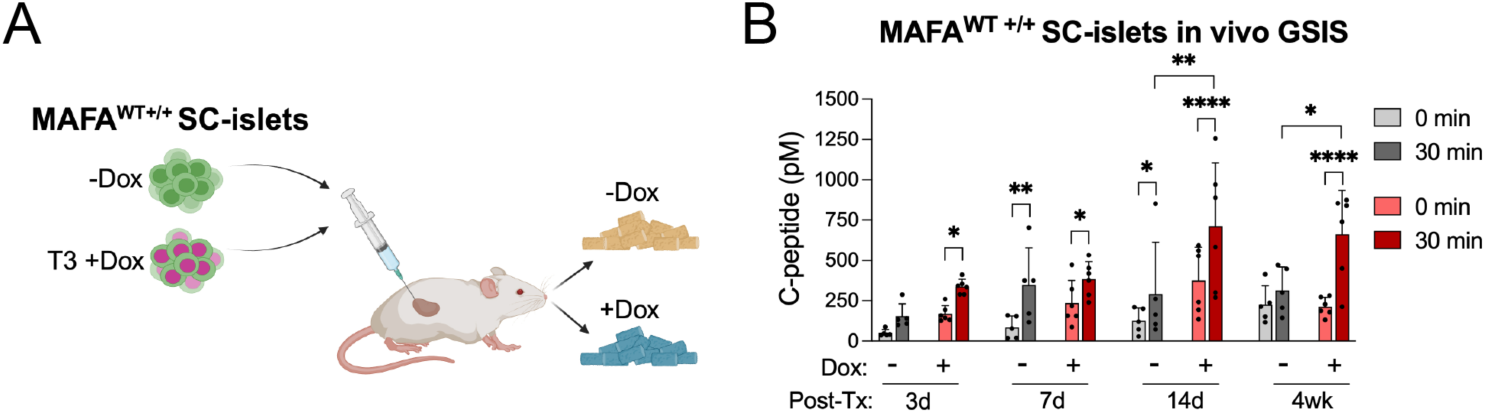
MAFA^WT+/+^ SC-islets exhibit improved functionality post-transplantation in mice. **(A)** Schematic showing MAFA^WT+/+^ SC-islets treated with (T3) or without dox to induce MAFA^WT+/+^ overexpression and transplanted under the kidney capsule of mice. Mice transplanted with MAFA^WT+/+^ T3 +Dox SC-islets were provided a doxycycline-containing diet. **(B)** Comparison of serum C-peptide levels of mice transplanted MAFA^WT+/+^ SC-islets -/+dox at matched post-transplantation time periods in the fasted state (0 min) or 30 min after glucose injection. p-values by two-way repeated measures Anova followed by Tukey’s multiple comparisons test. *p<0.05, **p<0.005, ***p<0.0005, ****p<0.00005. Data are presented as mean ± s.e.m (n≥3 biological replicates).

## Discussion

Despite the functional and transcriptional discrepancies between immature SC-islets and mature primary human islets when assessed *in vitro*^1–6,12,67^, ongoing clinical studies have demonstrated that several months post-transplantation, SC-islets mature in vivo and are capable of significantly reducing or abolishing the requirement of exogenous insulin for type 1 diabetes patients^8,68^. Acquisition of greater islet functionality is attributed in part to MAFA upregulation over time based on studies performed in primary human islets and SC-islet transplantation in mice^9,12^. Here, we use an inducible MAFA overexpression cassette to model MAFA-mediated functional maturation in SC-islets in vitro and in vivo. Simultaneously studying MAFA^WT^ and the MAFA^S64F^ MODY variant reveals important insight into biological processes that contribute to maintaining a heightened ability to secrete insulin. Identification of TFs and their target genes involved in these processes provides new avenues for improving SC-islet robustness and preventing dysfunction of endogenous islets.

A most striking contrast between MAFA^WT^ and MAFA^S64F^ overexpression was the timing at which they acquired functionality. While MAFA^WT^ overexpression gradually increased GSIS functionality over a two-week period, MAFA^S64F^ exhibited a comparable increase in GSIS within 72 hours before rendering the cells non-functional. Although short overexpression of MAFA^S64F^ increased mRNA expression for many of the same metabolic genes found in MAFA^WT^ cells, this did not translate to the protein level for many of the identified genes. Despite MAFA^WT^ and MAFA^S64F^ both being overexpressed using the same system, MAFA^S64F^ is more stable and expressed at higher levels. It is well documented that TF expression level (i.e. low versus high) can allow the execution of different transcriptional programs^69–71^. This suggests that timing and level of MAFA expression is essential to govern proper mature β cell identity and functional GRNs.

PC stood out as a protein that was highly upregulated in both MAFA^WT^ and MAFA^S64F^ at their most functional timepoints. PC is a TCA anaplerotic enzyme involved in increasing TCA flux by converting the final glycolytic metabolite pyruvate to oxaloacetate and replenishing TCA intermediates^24,25^. Consequently, both MAFA^WT^ and MAFA^S64F^ show increased pyruvate anaplerosis and increased mitochondrial respiration. These changes in glucose metabolism occurred at rates that mirrored GSIS functional acquisition in MAFA^WT^ and MAFA^S64F^ cells and are known to be critical for GSIS. However, while the changes in OxPhos metabolism occurred over many days in MAFA^WT+/+^ SC-islets, they occurred much more abruptly in MAFA^S64F^ SC-islets, potentially preventing glucose from serving as building block for biological processes that are essential for sustaining functionality and cellular homeostasis observed in MAFA^WT^ SC-islets.

Notable cellular processes that were upregulated in MAFA^WT^ SC-islets included Golgi organization, cilium organization, ribosomes, peroxisomes, PPP, fatty acid metabolism, circadian entrainment, glycolysis and OxPhos. Golgi organization, ribosomes, PPP, and fatty acid metabolism indicate a period of cellular growth to develop the cellular machinery required to accommodate increased metabolic demands, while cilium organization, peroxisomes, circadian entrainment, glycolysis and OxPhos point to a transition to a more sophisticated metabolic state with improved glucose responsiveness. MIMS-EM imaging demonstrated that MAFA^WT^ overexpression initially increases carbon and nitrogen retention within cells to be used as cellular building blocks and later switch to primarily oxidative metabolism of glucose and glutamine^72^, which is further supported by our Seahorse assays. Increased MAFA^WT+/+^ T3 mitochondrial oxidative metabolism coincides with enhanced GSIS functionality consistent with OxPhos being a key driver of islet glucose responsiveness. Conversely, MAFA^S64F+/-^ SC-islets rapidly upregulated oxidative metabolism and burn through metabolic fuels without ever retaining carbon or nitrogen for biogenesis purposes. Evidence that MAFA^S64F+/-^ SC-islets become non-functional and exhibit mitochondrial swelling point to metabolic activity overload as a possible mechanism for MAFA^S64F+/-^ dysfunction. Taken together, these studies indicate a holistic role for MAFA^WT^ in the development of β cells to accommodate for increased metabolic demands.

A key distinction between MAFA^WT^ and MAFA^S64F^ came from their inferred TF activity networks. Very early on, MAFA^WT+/+^ in T1 clusters was identified as the primary interconnected signaling hub indicative of MAFA being the main driver of downstream effects. Moreover, MAFA^WT+/+^ clusters at both T1 and T3 could be neatly grouped into communities based on shared target genes indicative of a coordinated progression towards cellular maturation. In contrast, MAFA^S64F+/-^ T1 cluster readily gave indication of a distressed and discordant TF activity network. The rapid increase in oxidative stress-associated TFs alongside findings from the bulk RNAseq and proteomics analysis indicating an upregulated response to oxidative stress and poor MAFA^S64F^ SC-islet survival when challenged with glucose oxidase indicated that MAFA^S64F^ T1 clusters were insufficiently equipped to handle the rapid rise in OxPhos metabolism. In contrast, MAFA^WT^ omics analysis combined with an oxidative stress challenge with glucose oxidase suggest that improved cellular stress handling is a vital role of MAFA during β cell maturation. As demonstrated by mouse transplantation studies, improving β cell resilience had direct implications for SC-islet functionality post-transplantation. Further investigation into the metabolic maturation of islets is expected to reveal targets for preventing β cell dysfunction and improving survival of functional β cells during periods of increased demand such as adolescence and high stress such as transplantation.

In summary, our study highlights the importance of MAFA for islet functional maturation. We identified MAFA-regulated processes that are important for metabolic maturation and homeostasis. Examining how cellular viability and function in response to cellular stress changes in more mature cells advances our understanding for how functional maturation impacts susceptibility to stress. Importantly, these studies are expected to more closely model a transitional period during β cell development rather than a fully matured adult state. Thus, the findings herein have implications for better harnessing biological processes to increase SC-islet functional maturation as well as the pathology of diabetes during adolescence.

## Author Contributions

Conceptualization, V.A.C., A.V.P., J.C., R.S., and M.H.; investigation, V.A.C., M.S., M.A.A., S.J., M.S.W., Y.L., A.B., J.M.-P., R.E.H., A.A., R.G. T.C.A., and R.AeD.; formal analysis, V.A.C., M.S., S.J., A.B., J.M.-P., T.C.A., and R.AeD; supervision, H.L., F.J.T., R.S., A.V.P., and M.H.; writing-original draft V.A.C.; writing-editing and revision V.A.C., A.V.P., and M.H.; funding acquisition, A.V.P., M.H., and R.S.

## Methods

### Stem cell culture and pancreatic differentiation

Mel1 INS^GFP/W^ ^73^human embryonic stem cells (hESCs) were maintained on mouse embryonic fibroblasts (MEFs) in hESC media composed of DMEM/F12, 20% v/v KnockOut serum replacement (Thermo Fisher Scientific), nonessential amino acids (Thermo Fisher Scientific), GlutaMAX (Thermo Fisher Scientific), 2-mercaptoethanol (Millipore) and supplemented with 4ng/ml recombinant human fibroblast growth factor 2 (FGF2; R&D Systems). Karyotyping was used to confirm the genomic integrity of hESC maintenance lines and mycoplasma testing was performed regularly to ensure that the cultures remained contamination free. Confluent hESCs were passaged by enzymatic dissociation into single cell suspension using TrypLE (Gibco) every 3-4 days.

To initiate differentiation, hESCs were enzymatically digested using TrypLE Select (Gibco) and 5.5x10^6^ cells were seeded in 5.5ml of hESC media supplemented with 10ng/ml of activin A (R&D Systems) and 10ng/ml of heregulinB (PeproTech) per well in a 6-well suspension plate. The plates were incubated at 37°C and 5% CO2 on an orbital shaker at 100 rpm to induce 3D sphere formation. After 24h, the spheres were differentiated using an adapted version of Parent et. al.^22^. Each well was collected into a 1.5ml Eppendorf tube and allowed to settle by gravity, washed once with RPMI (Gibco), and resuspended in d1 differentiation media in a 6-well suspension plate (5.5ml media/well). d0-d3 cells were fed by aspirating media from each well and replacing with 5.5ml of fresh media daily. d4-d20 spheres were fed using 5ml of media. Starting at d13, d12+ media (check media names) was supplemented with Aphidicolin (APH, Cayman Chemical). Cells were reaggregated at d19-20. Briefly, d19-20 spheres were collected into a 50ml falcon tube, rinsed with PBS, and dissociated using Accumax (STEMCELL Technologies) into a single cell suspension, strained through a 40 mm filter (Corning), and re-seeded at 4-5x10^6^ cells/well in D21+ media with 10 μM of the ROCK inhibitor Y-27632 (Tocris). The next day the media was replaced with D21+ media without Rock inhibitor Y-27632. Cells were cultured in D21+ media until d27-d30. Media compositions for differentiation of INS^GFP/W^ hESCs are as follows: d1, RPMI (Gibco) containing 0.2% FBS, 1:5,000 ITS (Gibco), 100 ng/mL activin A, and 1uM CHIR (STEMCELL Technologies); d2, RPMI containing 0.2% FBS, 1:2,000 ITS, and 100 ng/mL activin A; d3, RPMI containing 0.2% FBS, 1:1,000 ITS, 2.5 μM TGFbi IV (Calbiochem), and 25 ng/mL keratinocyte growth factor (KGF; R&D Systems); d4–5, RPMI containing 0.2% FBS, 1:1,000 ITS, and 25 ng/mL KGF; d6–7, DMEM (Gibco) with 25 mM glucose containing 1:100 B27 (Gibco) and 3 nM TTNPB (Sigma); d8, DMEM with 25 mM glucose containing 1:100 B27, 3 nM TTNPB, and 50 ng/mL epidermal growth factor (EGF; R&D Systems); d9–11, DMEM with 25 mM glucose containing 1:100 B27, 50 ng/mL EGF, and 50 ng/mL KGF; d12+, DMEM with 25 mM glucose containing 1:100 B27, 1:100 GlutaMAX (Gibco), 1:100 NEAA (Gibco), 10 μm ALKi II (Axxora), 500 nM LDN-193189 (Stemgent), 1 μm Xxi (Millipore), 1 μM T3 (Sigma-Aldrich), 0.5 mM vitamin C, 1mM APH (Cayman Chemical), 1 mM *N*-acetylcysteine (Sigma-Aldrich),10 μM zinc sulfate (Sigma-Aldrich), and 10 μg/mL heparin sulfate; d21+ Connaught Medical Research Laboratories (CMRL) medium containing 10% FBS, 1:100 Glutamax (Gibco), 1:100 NEAA (Gibco), 10 μm ALKi II (Axxora), 0.5 mM vitamin C, 1 μM T3 (Sigma-Aldrich), 1 mM N-acetyl Cysteine (Sigma-Aldrich), 10 μM zinc sulfate (Sigma-Aldrich), and 10 μg/ml of heparin sulfate.

### Generation of inducible MAFA expression clonal hESC lines

For temporal control of MAFA cDNA activity in INS^W/GFP^ hESCs, human MAFA cDNAs (Roland Stein) were subcloned into the AAVS1-TRE3G-EGFP (Addgene #52343) to generate iMAFA donor plasmids. TALENS targeting of iMAFA donor plasmids to INS^GFP/W^ AAVS1 using targeting plasmids using AAVS1-TALEN-L (Addgene #59025) and AAVS1-TALEN-R (Addgene #59026)^74^. 8x10^6^ cells in hESC media supplemented with FGF2, RI, and 100units/ml pencillin/streptomycin (P/S; Corning) were combined with 20 μg of iMAFA donor plasmid and 5μg of each targeting plasmid in a Gene Pulser Cuvette (Bio-Rad) and electroporated with the Gene Pulser Xcell System (Bio-Rad) using settings 250 V, 500 μF. Following electroporation cells were incubated in the cuvette on ice for 20 minutes before being plated on 10-cm plates with DR4 iMEFs (Thermo Fisher Scientific). Targeted cells were selected for using puromycin (5 mg/ml, Sigma-Aldrich), stocked and genomic DNA was extracted to verify correct insertion of iMAFA into the AAVS1 locus.

### Human islets

Human islets used in the experiments were obtained from the UCSF Islet Core and the Integrated Islet Distribution Program (IIDP). Following isolation and delivery, islets were handpicked under a stereomicroscope and cultured in PIM(S) media (Prodo Labs) in 10cm non-adhesive petri dishes (Thermo Fisher Scientific) for 24-72 hours prior to experiments to allow for recovery. All families of organ donors provided written consent for use in research. Islets were used with approval by the institutional biosafety committee at UCSF.

### Flow Cytometry

Spheroids were dissociated into single cell suspensions using Accumax, fixed in 4% paraformaldehyde (PFA; Electron Microscopy Sciences), permeabilized (Invitrogen™ eBioscience™ Permeabilization Buffer, Fisher Scientific) and stained for indicated cellular markers for analysis on BD LSRFortessa X20 Dual using FACSDiva software as described previously^4,75^. Data were analyzed with FlowJo software. Anti-human C-peptide and anti-glucagon were conjugated in-house using the Molecular Probes Antibody Labeling Kits according to the manufacturer’s instructions. Antibodies: PDX1 and NKX6.1 (BD Biosciences), C-peptide (Millipore), Glucagon (Sigma-Aldrich).

### Immunofluorescence staining

Spheroids were collected in 1.5ml Eppendorf tubes, allowed to settle by gravity and washed once with PBS before being fixed with 4% PFA for 15 min at room temperature. Samples were stored in PBS at 4 °C until embedding in Optimal Cutting Temperature (OCT; company) embedding. OCT embedded samples were frozen at -80 °C and sectioned at 20μm. For staining, samples were rinsed in PBS, blocked in CAS-block (Invitrogen) with 0.2% Triton X-100 (Fisher) for 1 hour at room temperature, and incubated with Insulin (Dako) and MAFA (Cell Signaling Technologies) primary antibodies diluted in CAS-block overnight at 4 °C. The next day, samples were washed in PBS containing 0.1% Tween-20 (PBS-T, Sigma) and incubated with Alexa Fluor (Thermo Fisher Scientific) secondary antibodies in CAS-block for 1 hour at room temperature. Samples were washed in PBS-T and mounted with coverslips using Prolong Diamond Antifade Mountant with DAPI (Thermo Fisher Scientific). Images were generated using a Nikon SP8 confocal microscope.

### Western Blotting

Spheroids were rinsed in PBS before being lysed in RIPA buffer (Thermo Fisher Scientific) supplemented with protease inhibitor cocktail and phosphatase inhibitor mini tablets (Thermo Fisher Scientific). Protein concentration was measured using DC Lowry assay (Bio-Rad), diluted to 1μg/μl in 4x Laemmli SDS sample buffer and boiled at 95 °C for 5 minutes. 20μg of cell lysates were resolved on 10% acrylamide SDS-PAGE gels (Bio-Rad) and then electro-transferred to PVDF membranes (Bio-Rad) and immunoblotted with MAFA (Cell Signaling Technologies) primary antibody in TBS with 0.1% Tween-20 (TBS-T) and 5% milk w/v. HRP-conjugated secondary antibodies (Jackson ImmunoResearch) were used and enhanced chemiluminescence detection using Supersign^TM^ West Femto Maximum Senstivity Substrate allowed for development using a ChemiDoc Touch (Bio-Rad). Blots were then stripped using Restore^TM^ Western Blot Stripping Buffer (Thermo Fisher Scientific) and re-probed for anti-TBP (Cell Signaling Technologies) as a loading control.. Unprocessed scans of western blots are shown.

### Dynamic GSIS Perifusion

Dynamic GSIS was performed using a PERI4 perifusion machine (BioRep). SC-islets were handPrior to dynamic GSIS, SC-islets were incubated in d21+ media with or without dox and counted using50 SC-islets or 20 human islets were loaded into chambers between two layers of polyacrylamide microbead slurry (BioRad). Under 37 °C temperature-controlled conditions clusters were perifused at 100μl/min in Krebs-Ringer Bicarbonate buffer (KRB) (115mM NaCl, 5mM KCl, 1mM MgCl2, 2.5mM CaCl2, 25mM HEPES, 24mM NaHCO3, 0.1% BSA). After an initial 1-2 hour preincubation in 2.8mM glucose-KRB, the outflow was collected every minute for the duration of the 80-minute protocol consisting of 20 minutes 2.8mM glucose-KRB, 30 minutes 16.7mM glucose-KRB, 20 minutes 2.8mM glucose-KRB, and 10 minutes of 30mM KCl in 2.8mM glucose-KRB. After the experiments, clusters were recovered from the chambers and lysed in acid ethanol. C-peptide levels were measured using the STELLUX Chemi Human C-peptide ELISA kit (ALPCO) and DNA was measured Quant-iT PicoGreen dsDNA Assay kit.

### Seahorse mitochondrial oxygen consumption assay

Real-time measurements of mitochondrial OCR were measured using XFe24 extracellular flux analyzer (Seahorse Bioscience). KRB buffer without NaHCO3 and pH adjusted (7.4) allowed for accurate measurements. 50 SC-islets and 20 human islets were washed in the appropriate KRB buffer containing 2.8mM glucose, placed in a well of an islet plate and secured with a mesh insert to prevent movement during the assay. Cluster-loaded plates were then incubated in a non-CO2 incubator at 37 °C° for at least 1 hour. Measurements were then taken at baseline 2.8mM glucose, 16.7mM glucose stimulation, oligomycin (5μM), carbonyl cyanide-4-(trifluoromethoxy) phenylhydrazone (FCCP, 1μM), and rotenone (5μM) and antimycin A (5μM) sequentially. At least two replicates were run per group. Clusters were collected at the end of experiments for C-peptide and DNA content. OCR was normalized to DNA content.

### Total RNA sequencing and analysis

Total RNA was extracted from SC-islets using RNeasy mini kit (Qiagen) according to manufacturer’s instructions. Genomic DNA contamination was removed using DNase I (RNase-free) (Qiagen). Samples were submitted to Novogene and those passing QC (RIN≥ 7.0) were used for downstream sequencing using the Novogene NGS RNA Library Prep Set (PT042) and the Illumina NovaSeq X Plus. Sequence alignment was performed against the human GRCh38/hg38 reference genome using HISAT2.

For quality control, counts were converted to transcripts per million (TPM) and samples were inspected by library size, PCA, hierarchical clustering and correlation heatmaps. Differential expression was analyzed with DESeq2^76^ using Wald tests after retaining genes with counts >10 in every sample included in the respective analysis. Log2 fold changes were shrunken with ashr^77^. Genes with an false discovery rate (FDR) threshold ≤ 0.05^78^ were considered significant; |log2 fold change| > 0.33 defined up- and downregulation. Over-representation analysis (ORA) for GO terms and KEGG pathways was performed using WebGestalt (WEB-based Gene SeT AnaLysis Toolkit). Statistical significance was determined using the Benjamini-Hochberg FDR correction, with a threshold set at FDR < 0.05.

### Proteomic Sample preparation

Islet samples were lysed in ice-chilled DOC lysis buffer (10 mM Tris-Cl pH 7.5, 1% sodiumdeoxycholate (SDC), 1 mM EDTA-Na, +1× complete) using a Precellys homogenizer (Bertin Technologies). Soluble proteins in the supernatant fraction were digested using the iST Kit (Preomics) according to the manufacturer’s protocol.

### Proteomics LC-MSMS measurements

The soluble fraction, dried peptides were reconstituted in 2% acetonitrile/0.1% trifluoracetic acid prior to LC-MSMS measurement in data-independent acquisition (DIA) mode on a Q-Exactive HF-X mass spectrometer (Thermo Scientific), online coupled to a nano-RSLC (Ultimate 3000 RSLC; Dionex). Tryptic peptides were trapped on a C18 pre column (300 µm inner diameter (ID) × 5 mm, Acclaim PepMap100 C18, 5 µm, 100 Å, LC Packings) at 30 µl/min flow rate followed by reversed-phase separation on a C18 analytical column (nanoEase MZ HSS T3 Column, 100 Å, 1.8 µm, 75 µm × 250 mm, Waters) at 250 nl/min flow rate in a 95 min nonlinear acetonitrile gradient from 3 to 40% in 0.1% formic acid. The precursor MS spectrum was acquired at a resolution of 120,000 (full width at half-maximum) with a mass range from 300 to 1650 m/z with automatic gain control target set to 3 × 10E6 and a maximum injection time of 120 ms. MSMS fragment spectra were recorded in 37 DIA windows with a variable width at a resolution of 30 000 and automatic gain control target set to 3 × 10E6. Normalized collision energy was set to 28, and the spectra were recorded in profile type.

### Quantitative proteomics data analysis

DIA raw files of the soluble fractions were processed in directDIA in Spectronaut 19 (Biognosys), applying BGS default settings with an additional filter for quantification on proteotypic peptides and using the Swissprot Human database (release 2020_02, 20432 sequences). Normalized protein quantifications were exported, and missing values were imputed from normal distribution with default settings in the Perseus software (MPI Martinsried, version 2.0.11.0). The average imputed normalized protein abundances for the individual treatment groups were used for calculation of abundance ratios and a Student’s T-test (with Benjamini-Hochberg correction) was used as hypothesis test within the Perseus software. Some samples were excluded from further analysis for quality reasons. The mass spectrometry proteomics data have been deposited to the ProteomeXchange Consortium via the PRIDE^79^ partner repository with the dataset identifier PXD083852. Functional enrichment analysis of GO and KEGG pathways was conducted using WebGestalt, applying a Benjamini-Hochberg adjusted *p*-value threshold of 0.05.

### Single Nucleus sample preparation and sequencing

To isolate nuclei, a low-input nucleus isolation protocol adapted from 10x Genomics was performed. Sorted cells were washed once with 1 ml PBS + 1% BSA, counted on the basis of Trypan blue staining and centrifuged. The cell pellet was resuspended in chilled lysis buffer (10 mM Tris-HCl (pH 7.4), 10 mM NaCl, 3 mM MgCl₂, 0.01% Tween-20, 0.01% Nonidet P40 Substitute, 0.005% Digitonin, 1% BSA, 1 mM DTT, and 1 U/µl RNase Inhibitor; 50 μl/sample) and placed on ice for 4 min. Samples were then washed with 500 μl of wash buffer and nuclei were centrifuged. To gradually change from wash to diluted nucleus buffer, cells were washed once in a 1:1 mixture of wash buffer and diluted nucleus buffer and subsequently once with pure diluted nucleus buffer. Isolated nuclei were then resuspended in 10 μl diluted nucleus buffer. Nuclei integrity and concentration were assessed by trypan blue staining. Samples were then immediately carried into transposition and GEM generation according to the Chromium Next GEM Single Cell Multiome ATAC + Gene Expression protocol (10x Genomics, CG000338), loading for a target recovery of 10,000 nuclei per sample.

ATAC and gene expression libraries were sequenced on an Illumina NovaSeq X+ platform following the recommendations from 10x Genomics at a target read depth of 35000 reads/nucleus. Raw reads from both library types were jointly aligned to the human GRCh38 reference built from Ensembl release 112 and pre-processed using the 10x Genomics CellRangerARC pipeline (7.1.0) for downstream analyses.

### Multi-omic sequence data processing and filtering

Gene expression matrices were generated with Cell Ranger and loaded per sample from filtered feature-barcode matrices; the RNA modality was extracted from each multiome object for independent processing. Eight samples were processed (VC001–VC007, VC009).

Doublets were identified per sample with SOLO. For each sample, a copy of the count matrix was filtered to genes detected in at least 10 cells, reduced to the 2,000 most highly variable genes (flavor=“seurat_v3”), and used to train an scVI model with default parameters. A SOLO classifier was initialized from the trained scVI model, and barcodes were called as doublets when assigned the doublet label and the difference between doublet and singlet scores exceeded 1.

Cells with fewer than 200 detected genes were removed. Mitochondrial (MT-), ribosomal (RPS/RPL), and hemoglobin (HB, excluding HBP) gene sets were annotated, and QC metrics computed with scanpy.pp.calculate_qc_metrics. Following the median absolute deviation (MAD) strategy of Heumos et al^80^., cells were flagged as outliers when any of log-transformed total counts, log-transformed number of detected genes, or the percentage of counts in the top 20 genes fell more than 5 MADs from the sample median. Cells exceeding 20% mitochondrial counts were flagged separately. Outlier and high-mitochondrial cells were removed within each sample.

Per-sample objects were concatenated, yielding 58,272 cells across 20,736 genes. Genes detected in fewer than 10 cells across the merged object were removed, retaining 17,895 genes. Raw counts were preserved in a dedicated layer, then counts were library-size normalized to 10,000 counts per cell and log1p-transformed

### snATAC quality control and peak processing

Accessibility quality control was performed per sample in Signac, using the CellRanger fragment files and Ensembl v112 annotation. For every nucleus we required the number of fragments in peaks, the TSS enrichment score and the nucleosome signal to fall inside sample-specific bounds, together with a fraction of fragments in peaks above 0.45–0.55, fewer than 0.1–0.2 % of cut sites in the ENCODE hg38 blacklist and fewer than 0.3 % mitochondrial reads (1 % for the lowest-quality sample). Thresholds were set individually per sample from the metric distributions because library complexity differed between samples; the full set is given in Table X. Only nuclei passing both modalities were kept, leaving 45,298 of 58,272 nuclei (77.7 %).

Because peaks are called per sample, the per-sample peak sets were merged into a common peak set across all eight samples, yielding 503,377 consensus peaks, and counts were re-quantified against this common set to produce a unified peak-by-cell matrix.

Peak accessibility was normalized by term frequency–inverse document frequency (TF-IDF), and peaks above the 50th percentile of total accessibility were selected as variable features (FindTopFeatures), retaining 251,985 peaks. Dimensionality reduction used latent semantic indexing; the first LSI component correlated with sequencing depth and detected peak number and was excluded. ATAC processing used Signac v1.13.0 and Seurat v5.0.3 under R v4.3.2, with muon v0.1.6 and mudata v0.2.3 in Python.

Nuclei passing RNA quality control (58,272) were intersected with nuclei having paired ATAC profiles and passing ATAC quality control (45,298). Moreover, one sample was lost due to a clog (VC009), yielding 41,638 nuclei for downstream analysis.

### DRVI latent representation, clustering, and cell-type annotation

Multiome data (41,638 cells) were loaded as a MuData object containing RNA, ATAC, and TF-activity modalities, following per-sample GEX. DRVI was trained on the RNA modality alone. Raw counts were restored to .X, library-size normalized (normalize_total) and log1p-transformed, and the 10,000 most highly variable genes were selected (scanpy.pp.highly_variable_genes, default flavour, subset=True) from 17,092 detected genes.

The model was set up with DRVI.setup_anndata on the unnormalized count layer (is_count_data=True), specifying sample of origin (Sample) as a categorical covariate to account for sample-level technical variation. DRVI was constructed with 128 latent dimensions, two hidden layers of 128 units in both encoder and decoder, and the default piecewise negative binomial likelihood, and trained for 400 epochs without early stopping with KL warm-up over the full schedule (n_epochs_kl_warmup=400). Training required ∼17 min on a single NVIDIA A100.

Latent representations were extracted with get_latent_representation(). A neighborhood graph was built directly on the 128-dimensional latent space (n_neighbors=10, use_rep=“X”) and embedded by UMAP (spread=1.0, min_dist=0.1, random_state=123). Clusters were called by Leiden community detection on this graph (resolution 0.7, n_iterations=2). Two clusters containing mixed genotype and timepoint composition were sub-clustered in a second pass (restrict_to, resolution 0.2, n_iterations=2), yielding 15 final clusters.

Cluster identities were assigned manually by joint inspection of (i) canonical hallmark marker expression, (ii) precomputed lineage signature scores (SC-EC, SC-β, SC-α, SC-δ, exocrine, mesenchyme, progenitor), (iii) cell-cycle phase assignment, and (iv) genotype and timepoint composition. Latent dimension statistics were computed with a vanished-dimension threshold of 0.5, and per-dimension gene scores were derived using both out-of-distribution and within-distribution interpretability approaches. Analyses used scanpy v1.11.5, anndata v0.12.7, and DRVI v0.2.3.

### Enhancer-driven gene regulatory networks

Regulatory networks were inferred with SCENIC+, using a cisTarget database, built specifically on the dataset’s 125,972 consensus peaks above the 75th accessibility percentile, scored against the SCENIC+ v10nr_clust motif collection.

Candidate enhancers were defined by cis-regulatory topic modelling in pycisTopic. Latent Dirichlet allocation models from 100 to 300 topics were compared by four model-fit metrics and the 300-topic model was selected. Topics were binarized by Otsu’s method and by their top 5,000 regions, and differentially accessible regions were computed per cell type, giving 614 region sets for motif enrichment.

Enhancer-driven gene regulatory networks were inferred with the SCENIC+ Snakemake pipeline. Each region set was tested with both cisTarget and DEM, and enriched motifs were assigned to transcription factors using direct and orthology-transferred annotations. TF-to-gene and region-to-gene links were scored by gradient-boosting regression, region-to-gene links additionally by Spearman correlation to assign a direction, and eRegulons were assembled by gene set enrichment analysis over regions within 150 kb of each gene, keeping eRegulons with at least five target genes. This produced 306 direct eRegulons (205 transcription factors, 10,567 target genes) and 213 extended eRegulons based on orthology-transferred motifs. Per-cell eRegulon activity was quantified with AUCell and used for cell-type specificity scoring.

GRN analysis used Scanpy 1.8.2, anndata 0.10.5, muon 0.1.6, pycisTopic 2.0a0, SCENIC+ 1.0a1 and Mallet 202108.

Differentially expressed genes (DEGs) for each condition were identified using a one-vs-rest Wilcoxon rank-sum test (Scanpy rank_genes_groups) comparing each cluster against all other clusters in the dataset, following library-size normalization and log-transformation. Genes were considered significantly upregulated in a given cluster if they met a Benjamini-Hochberg-adjusted p-value ≤0.05 and log fold-change ≥0.33.

Transcription factors (TFs) were validated for each condition using the following criteria: a regulon specificity score (RSS) reflecting cluster-specific activity; a minimum of 10 target genes with direct (non-extended), positive TF-to-gene correlation; the TF’s own expression meeting the same DEG significance criteria described above (adjusted p < 0.05, log fold-change > 0.33); and expression in at least 10% of cells within the cluster. Because SCENIC+’s default transcription factor reference list is derived from motif annotations aggregated across multiple external databases, and can therefore include genes without established sequence-specific DNA-binding activity, the resulting validated TF set for each condition was additionally cross-referenced against the curated human transcription factor catalog of Lambert et al. (2018), and any gene not present in this reference was excluded from downstream analysis.

### Signaling hub network analysis

For each cell cluster, transcription factor (TF)–TF co-regulatory networks were constructed from validated eRegulons (SCENIC+ v1.0a2^29^, direct positive regulons, ρ(TF2G) > 0, ≥10 target genes) whose target gene sets significantly overlapped that cluster’s differentially expressed genes (one-vs-rest Wilcoxon test via Scanpy v1.11.5^30^, log fold-change > 0.33, adjusted P < 0.05, expressed in ≥10% of cluster cells; hypergeometric overlap test with target genes, Benjamini-Hochberg FDR < 0.05, SciPy v1.15.2^81^ and statsmodels v0.14.6^82^). Nodes represent validated TFs; edges connect TF pairs with significant target-gene overlap, weighted by Jaccard similarity. Networks were constructed and analyzed using NetworkX v3.4.2^83^. Node importance was quantified as a composite centrality score, calculated as the mean of degree, betweenness, and eigenvector centrality, each min-max normalized within each cluster’s network prior to averaging to prevent any single measure (owing to differences in numeric scale) from dominating the composite score. Eigenvector centrality was computed separately for each connected component of the network to avoid numerical instability on disconnected graphs; TFs in components smaller than four nodes were assigned degree centrality as a substitute, as eigenvector centrality is not well-defined for very small components. All centrality measures were computed on the unweighted network topology. Hub concentration per cluster was summarized using the Gini coefficient of composite centrality scores across all validated TFs in that cluster’s network, with higher values indicating a more unequal, hub-dominated distribution of connectivity.

### Louvain Community Clustering Network Analysis

To identify transcription factor (TF) co-regulatory modules within each condition, we constructed a TF-TF specificity network from the set of validated, condition-specific TFs, in which each TF’s target-gene set was intersected with genes significantly upregulated in that condition. An edge was drawn between two TFs if the overlap between their target-gene sets was statistically significant by hypergeometric test (Benjamini-Hochberg FDR < 0.05), weighted by the Jaccard index of the shared target sets. Communities within this network were identified using the Louvain modularity-maximization algorithm^84^ (python-louvain), at a resolution selected independently for each condition as the widest contiguous range of resolution values over which the resulting partition remained stable (adjusted Rand index ≥ 0.9^85^ between adjacent resolution steps). Partition robustness was further confirmed by repeating community detection across 100 random seeds at the selected resolution and quantifying per-TF assignment consistency.

For each Louvain community identified within a condition’s TF specificity network, a representative gene set was defined as the union, across all member TFs, of their validated direct target genes intersected with that condition’s significantly upregulated gene set (as defined above). Gene Ontology Biological Process and KEGG pathway enrichment for each community’s gene set was performed using g:Profiler, with a custom background consisting of all genes tested in the corresponding one-vs-rest differential expression comparison (rather than g:Profiler’s default whole-genome background), and significance assessed using g:Profiler’s FDR-based method (adjusted p < 0.05). Redundant enriched terms — defined as term pairs with Jaccard gene-overlap similarity ≥ 0.5 to a more significant, already-retained term — were collapsed to their single most significant representative.

### Steady State ^13^C-labeling studies

SC-islets were initially subjected to a 2-hour pre-incubation period in DMEM media (Sigma-Aldrich) supplemented with 10mM HEPES, 0.5% BSA (A7030, Sigma-Aldrich) 11mM Glucose (Sigma-Aldrich) and 2mM glutamine (Sigma-Aldrich). After 2 hours, media was replaced with fresh media of similar composition containing ^13^C-glucose and ^15^N-glutamine tracers and incubated for an additional 3 hours. For MIMS-EM studies, SC-islets were simultaneously incubated with ^13^C-glucose and ^15^N-glutamine. For Metabolic tracing studies, SC-islets were divided into three groups and each group was incubated with a metabolic tracer: [4-^13^C]glucose, and [5-^13^C]glucose. All isotope tracers were obtained from Cambridge Isotope Laboratories.

### Sample preparation for Correlative Electron Microscopy and Multi-isotope Mass Spectroscopy (MIMS-EM)

MIMS-EM experiments were performed as previously established by us^63,65^. Here, after steady state labeling with ^13^C-glucose ^15^N-glutamine tracers, SC-islets were then rinsed in PBS and stored 4°C in fresh fixative (2% formaldehyde and 2.5% glutaraldehyde in 0.15M sodium cacodylate buffer with 2mM CaCl2). Next, each SC-islet organoid sample was washed for 1 hour at room temperature using 0.15 M cacodylate buffer and then post-fixed in 2% osmium tetroxide and 1.5% potassium ferrocyanide solution in 0.15 M sodium cacodylate buffer. Samples were washed in double distilled water (ddH20) and incubated in 0.5% thiocarbohydrazide for 30 minutes followed by five ddH20 washes at room temperature. SC-islet were then incubated in 2% aqueous osmium tetroxide solution for 1 hour followed by extensive ddH20 washes. Next, they were transferred to a 2% aqueous uranyl acetate solution at 4 °C overnight. Samples were washed with ddH20 and placed into Walton’s lead aspartate solution prior to baking for 30 min at 60 °C using a bench-top baking oven. Baked SC-islet samples were washed with ddH20 and serially dehydrated using ice-cold ethanol at 70%, 90%, and 100% EtOH followed by dry acetone (10 minutes each step on ice). Dehydrated material was placed into 1:3, 1:1, and 3:1 solutions of Durcupan ACM:acetone for 12 hours in each concentration for sample embedding. Embedded samples were exposed to three changes of 100% Durcupan ACM for 24 hours each before being baked for 48 hours at 65 °C for solidification.

### MIMS-EM image acquisition and data analysis

To image whole cross-sections of SC-islets, we created large field-of-view (FoV) maps of SC-islet preps. Here, 80 nm-thick sections were cut using an ultramicrotome (Leica UC7) and arranged on 10 x 10 mm silicon wafers (University Wafers) and mapped using scanning electron microscopy (SEM) on a Zeiss Crossbeam 550 microscope (Zeiss, Germany). Initial user-supervised image acquisition was performed to guide automated tile acquisition and image mosaicking (Atlas 5, Fibics, Ottawa, Canada). Images were acquired with a pixel size of 5 nm and covered areas of approximately 2 SC-islets per sample. Next, sample wafers were transferred to a multi-isotope mass spectrometry microscope (MIMS) (50L NanoSIMS, Cameca, France) for acquisition of isotope maps (^13^C, ^12^C, ^32^S, ^15^N, ^14^N, and 31P). The following image acquisition parameters were used: image size of 512×512 pixels, raster size of 30-to-40um2, at least three frames per raster with a 10 min acquisition time per frame using the beam adaptor D1-3.

### Metabolic tracing

Following steady state labeling with one of three metabolic tracers described above, SC-islets were washed with ice-cold PBS, immediately frozen, and stored at -80°C until shipped for further analysis. Metabolites were extracted with quenching solution (20% Methanol (HPLC grade), 0.1% Formic acid, 3mM NaF, 100uM EDTA). Cellular extracts were frozen and freeze-dried (Labconco) before preparation for MS analysis. Dried extracts were resuspended in water containing 10µM D4-Taurine used as external loading control (Cambridge Isotope Laboratories). Samples were kept in an autosampler at 4 Celsius before being injected onto a Hypercarb column (100x4.6mm, 3μm, ThermoFisher) and eluted with a solution containing 7mM ammonium formate, 10μM EDTA and 2% acetonitrile. Ionization by electrospray took place in a SCIEX 7500 QTRAP. The Multiple Reaction Monitoring (MRM) masses used to detect lactate, glutamate and malate were 89/43, 90/43, 90/44 (lactate), 146/102, 147/102, 147/103, 148/103, 148/104 (glutamate) and 133/115, 134/116, 135/117 (malate). MS data was integrated using El-Maven software (v0.12.0) and the ensuing areas were used to calculate ^13^C-enrichments (AreaMRM/AreaTotal for each metabolite). Enrichments were also calculated for unlabeled standards and used to correct for natural abundance (defined as Atom percent excess, APE). The fractional contribution of pyruvate dehydrogenase to the synthesis of acetyl-CoA (*fPDH*) was calculated from the [5-^13^C]glucose-labeling experiments (Eqs. 1, 2). An approximation to the fractional contribution of pyruvate carboxylase to the synthesis of OAA (*fPC*) was calculated from the [4-^13^C]glucose-labeling experiments (Eq. 3).

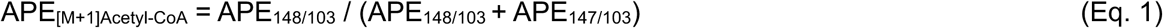

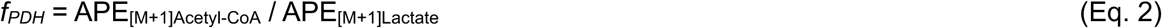

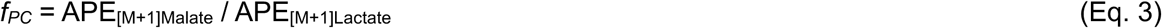

### Live-cell imaging

*In vitro* assessment of SC-islet resilience to oxidative stress was performed on an Incucyte Live-Cell Imaging System (Sartorious). SC-islets were added to a 96wp (Corning) in 200ul of d21+ media. 10mU/ml glucose oxidase (Sigma Aldrich) was added to treatment wells to induce H2O2-mediated oxidative stress. Brightfield and GFP images were acquired every hour over a 24-hour treatment period. Viability of healthy insulin-expressing SC-islets was quantified as integrated GFP signal (GCU μm^2^/image) using Incucyte Base Analysis Software (Sartorious). GFP signal was normalized as a percentage of maximal GFP signal (time point 0) and GOx-treated wells were further normalized to genotype-matched untreated control GFP at each time point.

### Mouse transplantation studies

NOD.Cg-*Prkdc^scid^ Il2rg^tm1Wjl^* Tg(Ins2-HBEGF)6832Ugfm/SzJ (NSG.RIP-DTR) mice were obtained from Jackson Laboratories and bred at UCSF for these studies. SC-islet clusters were transplanted under the kidney capsule of mice (3-5x10^6^ cells per mouse) according to protocols approved by the UCSF Institutional Animal Care and Use Committee. Mice transplanted with MAFA^WT+/+^ T3 SC-islets were fed a doxycycline-containing diet (200mg/kg, BioServ) while control mice were fed a normal diet. For in vivo glucose challenge experiments, transplanted mice were fasted for 5h followed by blood collection before and 30 minutes after an intraperitoneal injection of D-glucose solution (2g/kg). Human c-peptide in plasma samples was quantified by ELISA (Alpco #80-CPTHU-CH05).

