## Supplementary material for "MAFA drives pancreatic islet maturation through coordination of cell growth, metabolic maturation, and resilience to stress": Supplmental Figures

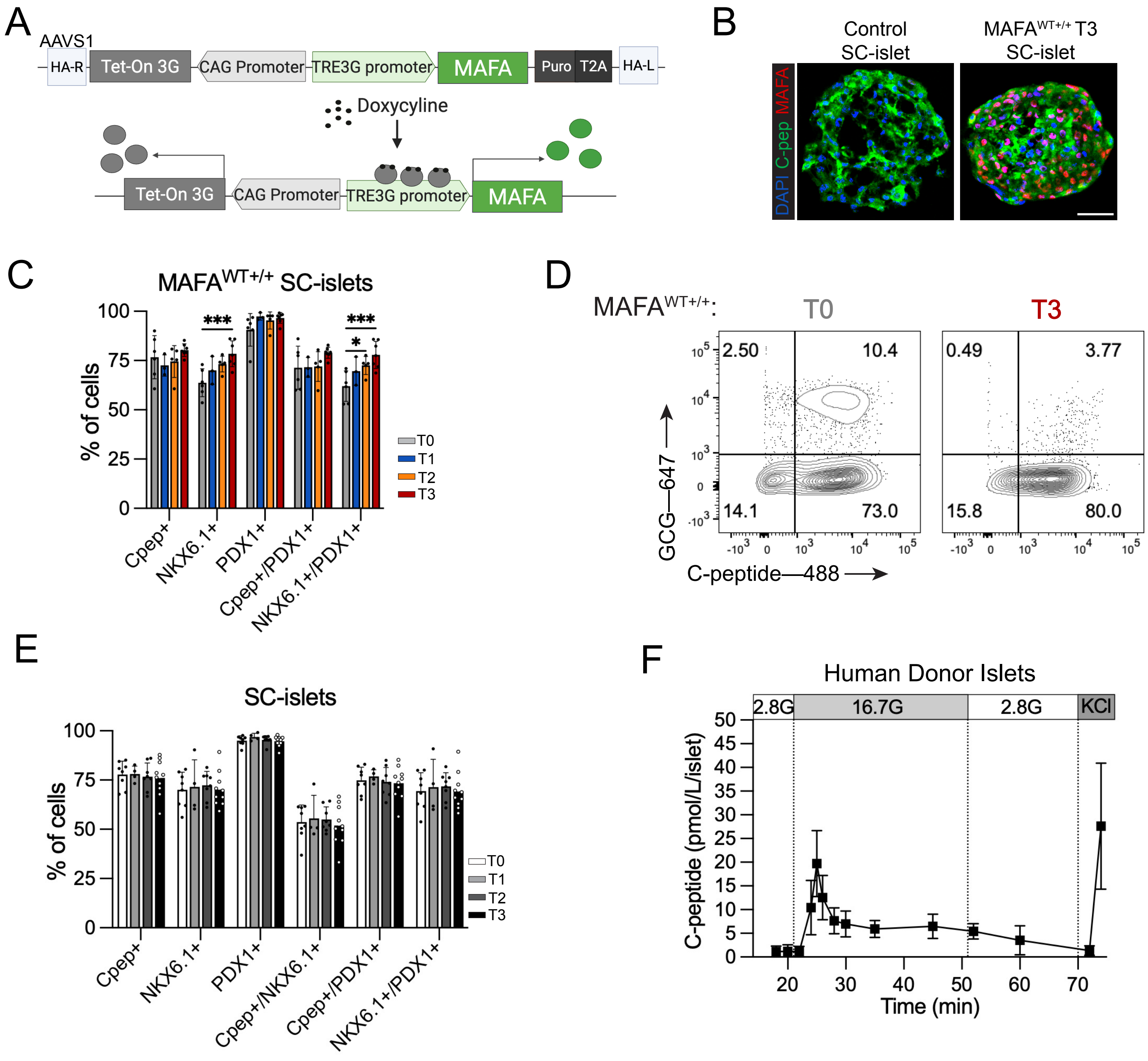


**Supplemental Figure S1**

**Supplemental Figure S1.** **(A)** Schematic of MAFA doxycycline-inducible overexpression construct used in combination with TALENs approach to target the AAVS1 locus. **(B)** Immunofluorescence images for MAFA and C-peptide in SC-islets negative (left) or positive (right) for the MAFA overexpression cassette and exposed to doxycycline. Scale bar 50$\mu$m. **(C)** Flow cytometry quantification for each $\beta$ cell identity markers in MAFA$^{WT+/+}$ SC-islets. p-values were determined by two-way Anova followed by a Dunnett's multiple comparison versus non-overexpressing controls. **(D)** Flow cytometry of MAFA$^{WT+/+}$ SC-islets for GCG and C-peptide. **(E)** Flow cytometry quantification of $\beta$ cell identity markers in SC-islets negative for the MAFA overexpression cassette and exposed to dox over the course of the differentiation protocol. Statistical analysis was performed using two-way Anova and a Dunnett's multiple comparison versus SC-islets not exposed to dox. **(F)** Human donor islets GSIS assay normalized for C-peptide (pmol/L) normalized per islet.

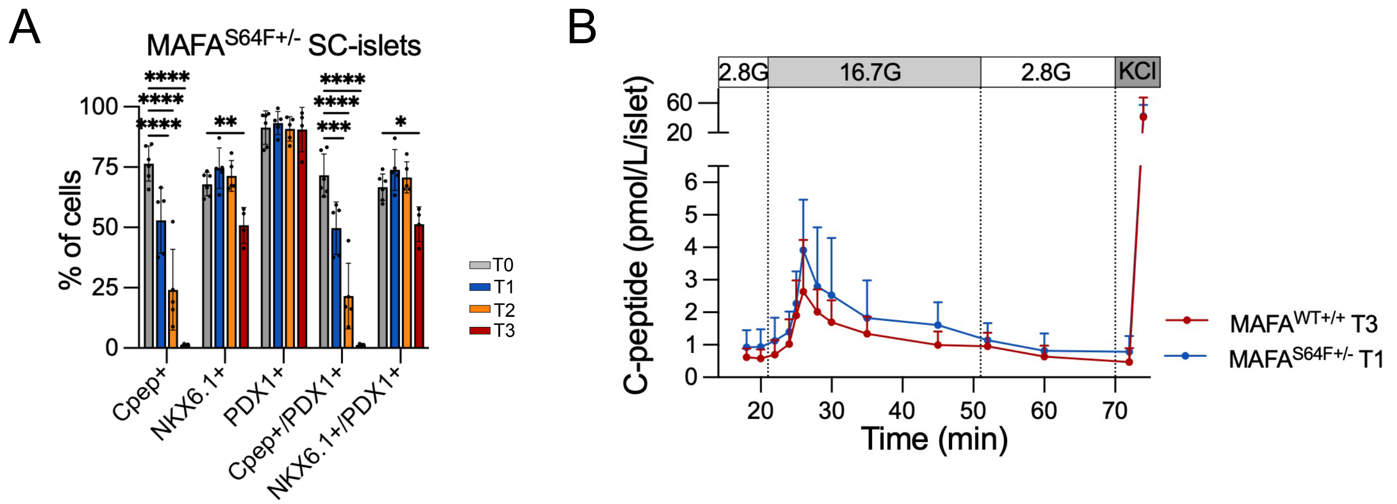


**Supplemental Figure S2**

**Supplemental Figure S2. (A)** Flow cytometry quantification for each β cell identity markers in $MAFA^{S64F+/-}$ SC-islets. p-values were determined by two-way Anova followed by a Dunnett's multiple comparison test versus non-overexpressing controls. **(B)** Comparison of $MAFA^{S64F+/-}$ T1 and $MAFA^{WT+/+}$ T3 SC-islet GSIS functionality. p-values by two-way repeated measures Anova followed by Šídák's multiple comparisons test.

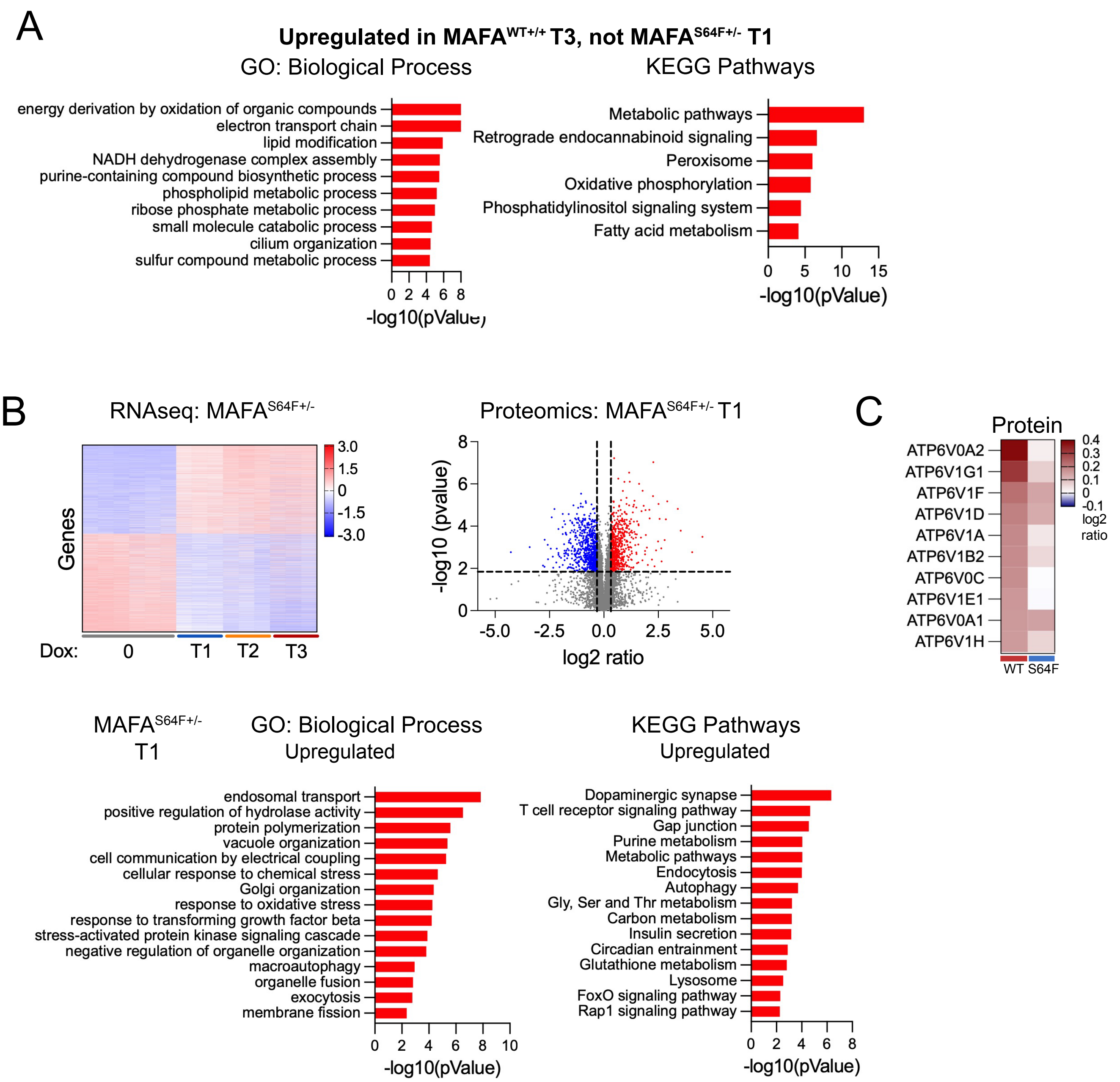


### Supplemental Figure S3

**Supplemental Figure S3. (A)** Functional analysis of $MAFA^{WT+/+}$ genes and proteins upregulated at T3 that are not upregulated in $MAFA^{S64F+/-}$ T1 cells. **(B)** Functional analysis of genes and proteins significantly downregulated in $MAFA^{WT+/+}$ T3 SC-islets. **(C)** V-ATPase protein expression in $MAFA^{WT+/+}$ T3 and $MAFA^{S64F+/-}$ T1 SC-islets.

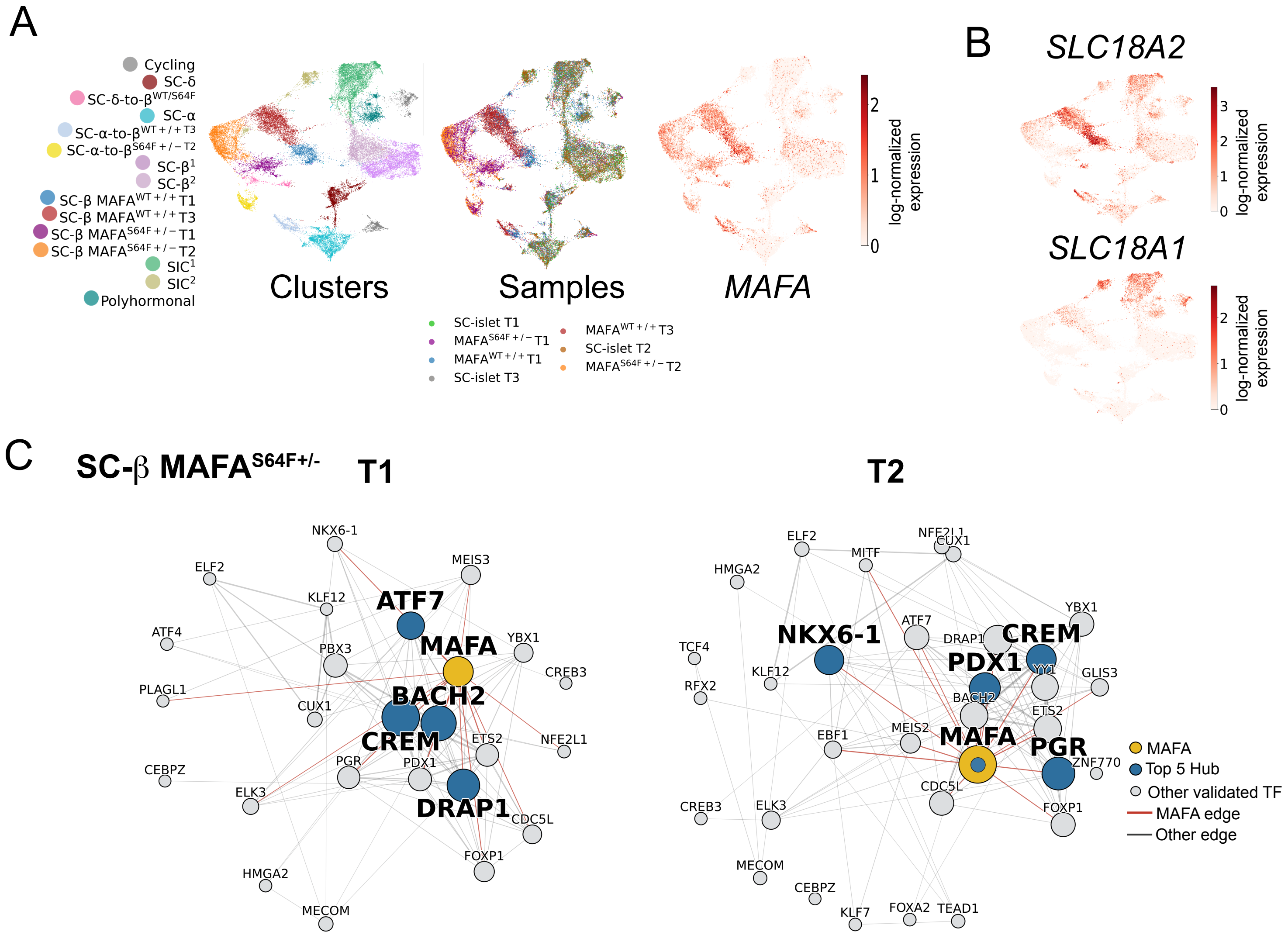


### Supplemental Figure S4

**Supplemental Figure S4. (A)** Multi-omics UMAPs showing clusters, samples, and *MAFA* expression. **(B)** UMAP expression of the *SLC18A2* and *SLC18A1* monoamine transporters. **(C)** Hub signaling networks for $MAFA^{S64F+/-}$ T1 and T2 clusters.

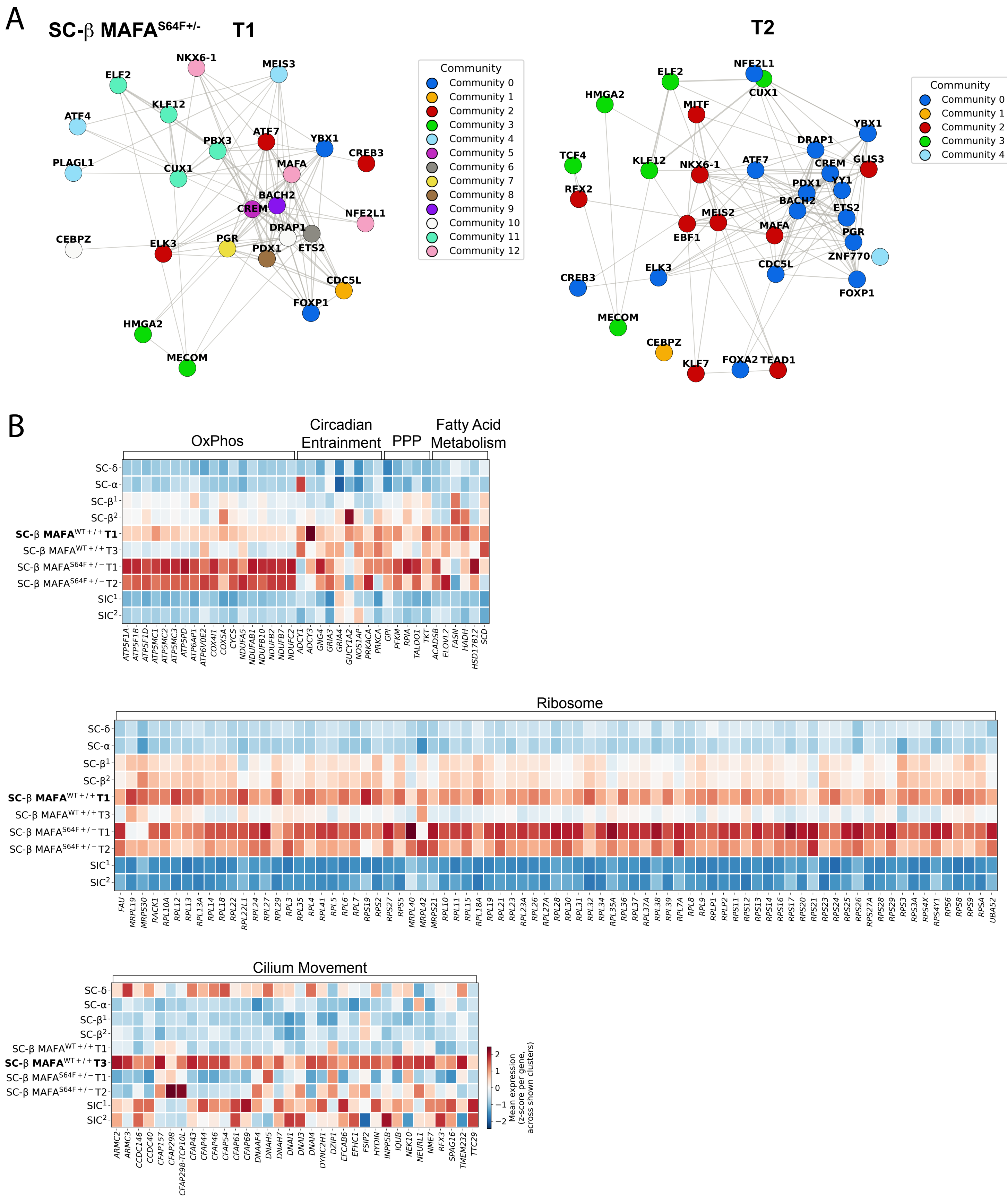


### Supplemental Figure S5

**Supplemental Figure S5. (A)** Louvain community clustering for MAFA$^{S64F+/-}$ T1 (resolution 2.68) and T2 (resolution 0.95) clusters. **(B)** snRNAseq heatmap expression of selected GO and KEGG pathways.

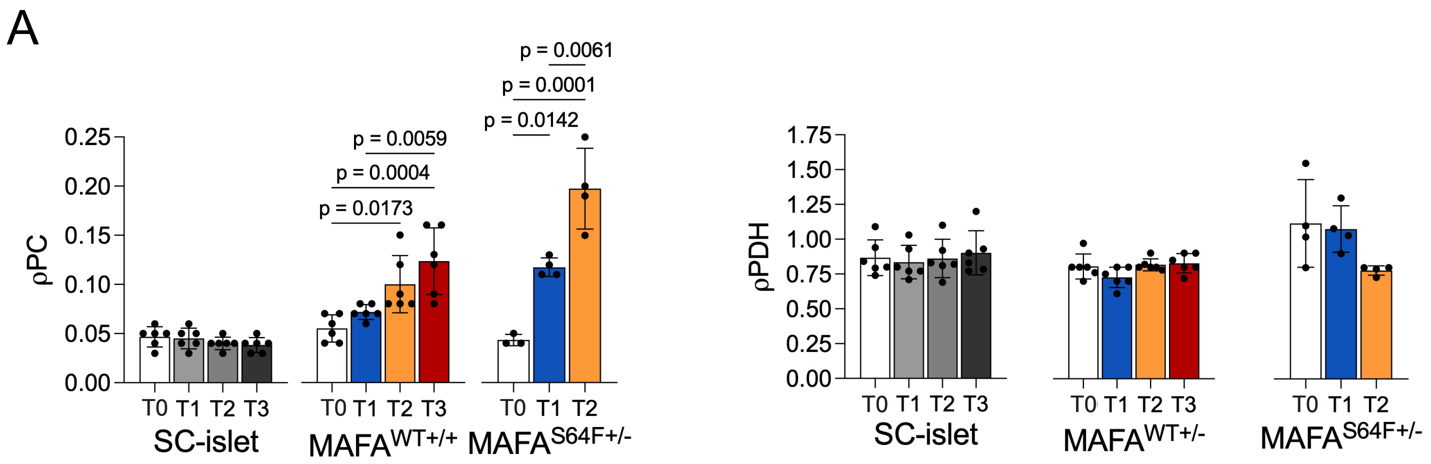


### Supplemental Figure S6

**Supplemental Figure S6. (A)** Metabolic tracing experiments for PC flux and PDH flux.

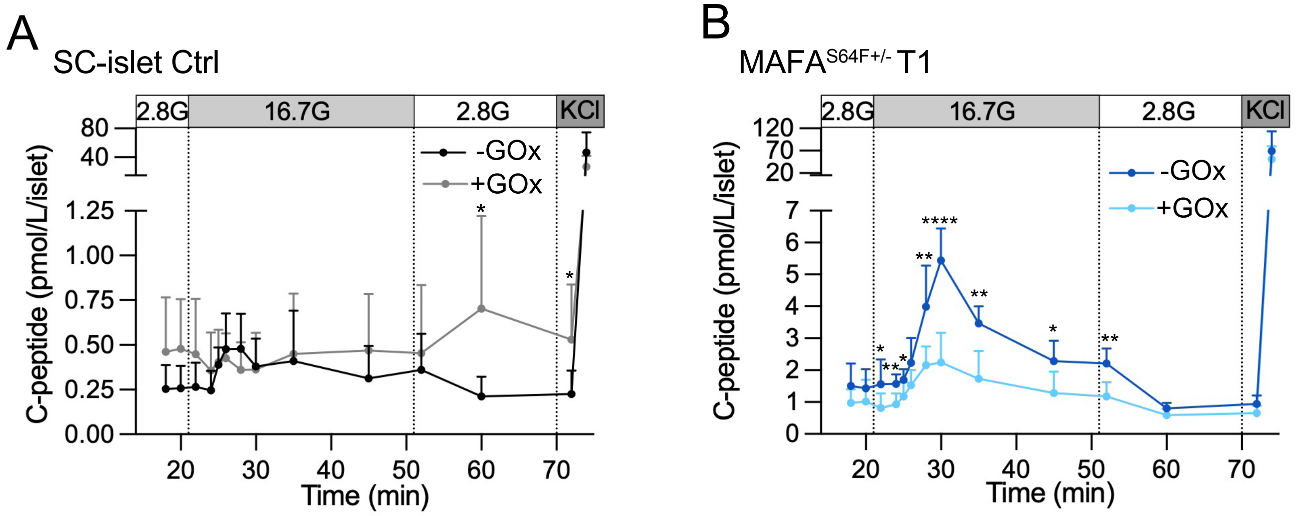


**Supplemental Figure S7**

**Supplemental Figure S7. (A)** Control SC-islets treated with glucose oxidase (10 mU/ml) to induce sustained oxidative stress and assessed for GSIS. p-values by multiple unpaired t-tests, with the false discovery rate controlled using the two-stage step-up method of Benjamini, Krieger, and Yekutieli ($q < 0.01$). **(B)** $MAFA^{S64F+/-}$ T1 SC-islets treated with glucose oxidase (10 mU/ml) to induce sustained oxidative stress and assessed for GSIS. p-values by multiple unpaired t-tests, with the false discovery rate controlled using the two-stage step-up method of Benjamini, Krieger, and Yekutieli ($q < 0.01$). *$p<0.05$, **$p<0.005$, ***$p<0.0005$, ****$p<0.00005$
